# The virulence of the *Cryptococcus neoformans* VNIa-5 lineage is highly plastic and associated with isolate background

**DOI:** 10.1101/2020.02.24.962134

**Authors:** Trieu Phan Hai, Thanh Lam Tuan, Duong Van Anh, Trinh Nguyen Mai, Lan Nguyen Phu Huong, Guy E. Thwaites, Errin Johnson, Nguyen Van Vinh Chau, Stephen Baker, Philip M. Ashton, Jeremy N. Day

**Affiliations:** Oxford University Clinical Research Unit, Wellcome Trust Asia Africa Programme, 764 Vo Van Kiet, Quan 5, Ho Chi Minh City, Vietnam; Hospital for Tropical Diseases, 764 Vo Van Kiet, Ho Chi Minh City, Viet Nam; Nuffield Department of Clinical Medicine, University of Oxford, Old Road Campus, Headington, Oxford OX3 7BN UK; Sir William Dunn School of Pathology, University of Oxford, S Parks Rd, Oxford OX1 3RE, United Kingdom UK; Cambridge Institute of Therapeutic immunology and Infectious Disease, Department of Medicine, University of Cambridge, Cambridge, UK

**Keywords:** Cryptococcus neoformans, Meningitis, Immunocompetent, HIV, Virulence, Pathogenesis

## Abstract

*Cryptococcus neoformans* most frequently causes disease in immunocompromised patients. However, in Vietnam and east Asia, disease is frequently reported in apparently immunocompetent patients. We have previously shown that almost all such disease is due to a specific lineage of *C. neoformans* – VNIa-5. However, in HIV-infected patients, infections due to this lineage are not associated with worse outcomes. Here, we demonstrate that the VNIa-5 lineage presents different virulence phenotypes depending on its source. Isolates derived from immunocompetent patients are more virulent than those from HIV-infected patients or the environment. Moreover, the virulence phenotype is plastic – sterile culture filtrate from highly virulent VNIa-5 strains can induce increased virulence in less virulent VNIa-5 isolates, which in turn can then induce increased virulence in their low virulence states. We present evidence that this phenomenon is driven by secreted proteins associated with extra-cellular vesicles.

## Introduction

Cryptococcal meningitis is a devastating disease due to infection with encapsulated yeasts of the *Cryptococcus* genus. The vast majority occurs in HIV-infected patients due to infection with *Cryptococcus neoformans*, and has a high moratlity rate ^1^. Disease in HIV-infected patients has been driven by the clonal expansion of a small number of well-defined lineages. These lineages are widely dispersed globally, but a single lineage predominates in most countries^2^. However, Vietnam is atypical and has two co-dominant circulating lineages (VNIa-5 and VNIa-4), each accounting for approximately 35-40% of cases of meningitis in HIV-infected patients^2-4^.

In addition to HIV-associated disease, cryptococcal meningitis is also well-described in HIV-uninfected patients in Southeast and East Asia^5-9^. Such patients account for approximately 20% of cases at our hospital in Vietnam^9^; the majority of these patients are apparently immunocompetent. Outcomes are similar to those in HIV patients, with 3 month mortality rates in the order of 20-30%^9^. Contrary to other locations in tropical and sub-tropical areas^10,11^, most infections in these patients (80%) are due to *C. neoformans*, rather than *C. gattii*^12^. We previously identified an association between the *C. neoformans* VNIa-5 lineage and disease in these apparently immunocompetent patients, it accounting for approximately 90% of meningitis due to *C. neoformans*^2,3,12^. Furthermore, HIV-uninfected patients with cryptococcal meningitis due to other lineages were significantly more likely to have co-morbidities associated with immune compromise^12^. These data suggest that VNIa-5 isolates have increased pathogenic potential. Of note, we do not observe clustering of isolates within the VNIa-5 lineage according to host immune status, signifying that the entire lineage has the potential to cause disease in immunocompetent people^2^.

Here, we aimed to understand the pathogenic ability of isolates of the VNIa-5 lineage. Using the *Galleria mellonella* infection model, we identified differences in virulence between isolates of the VNIa-5 lineage of different ecological backgrounds. The most virulent isolates were from HIV-uninfected patients, which were more virulent than VNIa-5 isolates from HIV-infected patients or the environment. However, we could induce increased virulence in lower virulence VNIa-5 isolates via transfer of sterile culture filtrate derived from highly virulent VNIa-5 isolates from immunocompetent patients. Consequently, these ‘induced’ isolates could sequentially increase the virulence of other low virulence isolates, and their own ‘naïve’ self. This virulence plasticity is mediated by peptide/protein associated with extracellular vesicles and likely underlies the ability of the VNIa-5 lineage to cause disease in immunocompetent people.

## Results

### Assessing the virulence of C. neoformans VNIa-5

We exploited the *Galleria* model of infection to compare the relative virulence of 20 VNIa-5 clinical isolates from HIV-uninfected patients with 20 VNIa-4 isolates from HIV-infected patients (Table S1). VNIa-5 infected *Galleria* had a significantly increased hazard of death compared with VNIa-4 infected *Galleria* (Hazard Ratio (HR) 1.4, 95% Confidence Interval (95CI) 1.2-1.6, P<0.001) (Figure 1A). After comparing the relative virulence of VNIa-5 isolates according to their source we found that *Galleria* infection with isolates from immunocompetent patients was associated with a significantly increased hazard of death compared with infection with isolates from HIV-infected patients (HR 2.2. 95CI 1.6 – 3.0, P<0.001), or with isolates from the environment (HR 5.7, 95CI 3.9 – 8.4, P<0.001, Figure 1B).

**Figure 1A:**
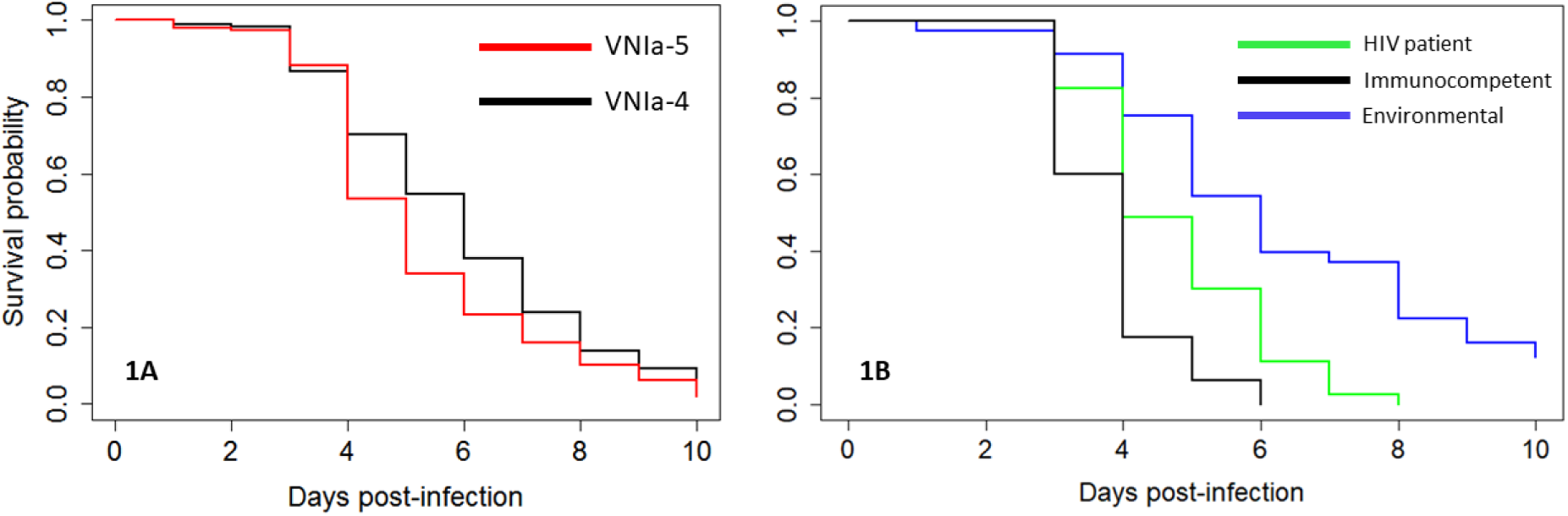
Survival curves of *Galleria mellonella* infected with human cerebrospinal fluid derived isolates of *Cryptococcus neoformans* of lineage VNIa-5 (derived from immunocompetent patients) or lineage VNIa-4 (from HIV-infected patients). N = 600 (20 isolates of each lineage, 15 larvae infected with each isolate, Hazard Ratio (95% Confidence interval): HR 1.4, 95CI 1.2,1.6; P<0.001 VNIa-5 versus VNIa-4). **Figure 1B: Survival curves of *G. mellonella* infected with *C. neoformans* lineage VNIa-5 isolates derived from immunocompetent patients (black line), HIV patients (green line), or the environment (blue line)**. 2 isolates of each source, 40 larvae per isolate. Hazard ratios and 95%CI for death: HIV derived vs environmental: 2.6, 95CI 1.8, 3.7; P< 0.001; immunocompetent derived versus environmental: 5.7; 95CI 3.9, 8.4; P< 0.001; Immunocompetent versus HIV derived: 1.7; 95CI 1.3 – 2.4, P<0.001. N=80 per arm.

Hypothesising that the difference in virulence phenotype by ecological background was a function of a previous infection experience we six-fold passaged environmental organisms through *Galleria*. The virulence phenotype of the environmental isolates remained stable over these multiple passages through *Galleria* with no change in the hazard of death between the infection with the ‘naïve’ environmental isolate versus infection with that isolate following passage (Figure S1).

We compared the expression of *in vitro* phenotypic characteristics associated with virulence in VNIa-5 isolates depending on their source (immunocompetent patients or HIV-infected patients). We compared 15 isolates from immunocompetent patients with 15 isolates from HIV-infected patients and measured growth rates in YPD broth and pooled human cerebrospinal fluid (CSF), capsule size, extracellular urease, phospholipase, and laccase activity. There was no difference in cell diameter, melanin production, urease, or phospholipase production. Isolates from HIV-uninfected patients had moderate but significant increased growth in YPD by 48 hours at 30°C and 37°C compared with isolates from HIV-infected patients (median 4.5×10^5^ CFU/ml (Interqartile Range (IQR) 3.6 X10^5^ - 5.2 ×10^5^) and 4 ×10^5^ CFU/ml (IQR 2.6 ×10^5^ - 5 ×10^5^) for immunocompetent isolates versus 3.3 ×10^5^ CFU/ml (IQR 2.9 ×10^5^ - 4.5 ×10^5^) and 2.8 ×10^5^ CFU/ml (IQR 2.2 ×10^5^ - 3.8 ×10^5^) for HIV derived isolates, P=0.004 and 0.002, respectively). The difference in the ability of isolates derived immunocompetent patients to grow at 37°C compared with 30°C did not reach statistical significance(P=0.058); however, there was statistically significant reduction in growth of isolates from HIV-infected patients at 37°C (P = 0.044) (Figure 2).

**Figure 2:**
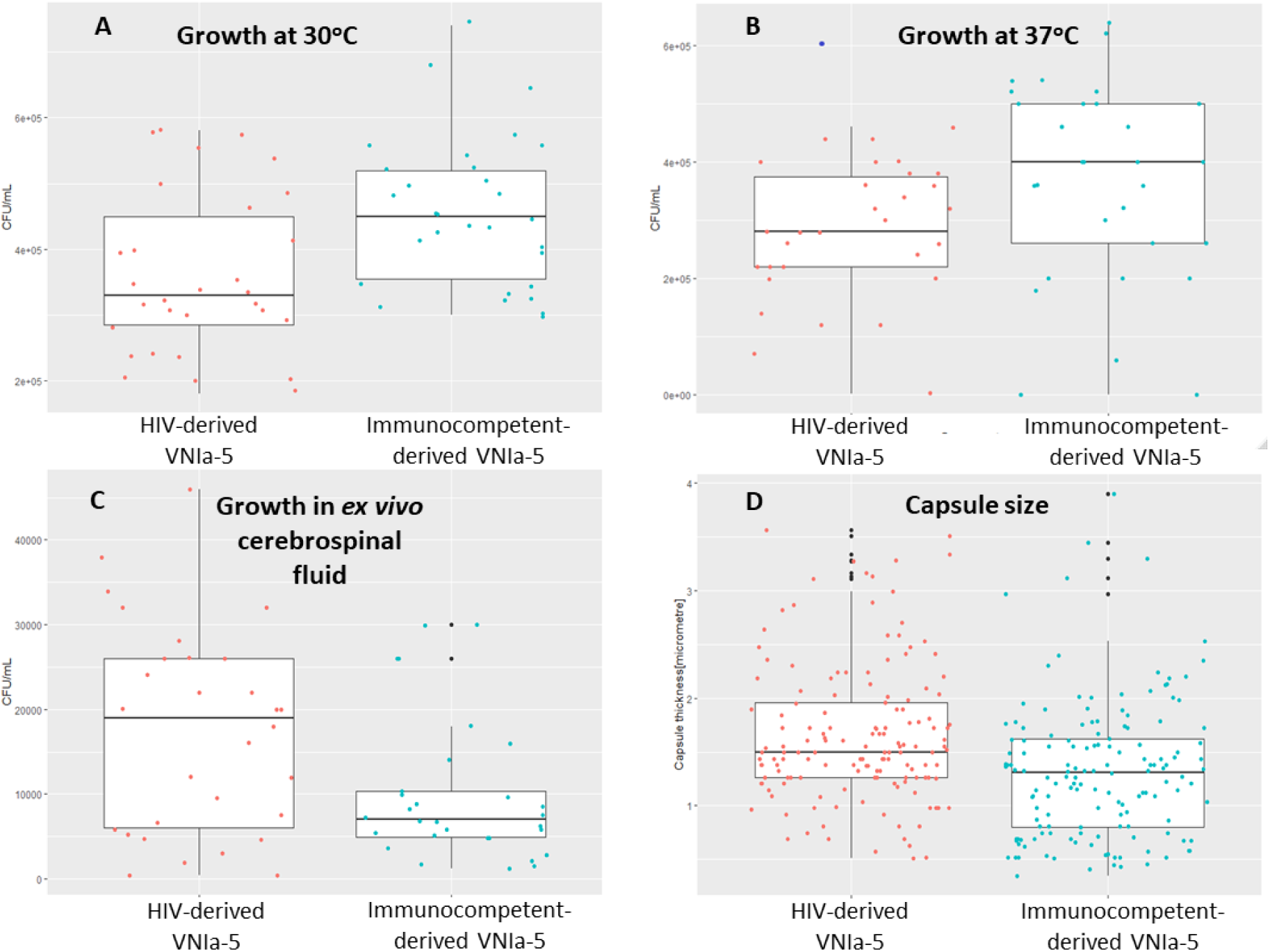
Comparative growth rates of VNIa-5 isolates in YNB broth at 30°C (panel A), 37°C (Panel B) an in *ex vivo* cerebrospinal fluid (C), and differences in capsule size *in vitro* (D) according to source – HIV-infected patients or immunocompetent patients. There was statistically significant greater growth of isolates derived from immunocompetent patients at both temperatures (P=0.004 and P=0.002 for 30°C and 37°C respectively). There was reduced growth at 37°C for both sets of isolates, but this was only statistically significant for those derived from HIV-infected patients. In contrast, isolates derived from immunocompetent patients appeared to have significantly impaired growth in *ex vivo* CSF compared with isolates from HIV-infected patients (P=0.02). (Note that the Y-axis scales are not the same in each panel). For all experiments N=15 isolates of each type, with 2 biological replicates.

In contrast, isolates derived from immunocompetent patients had slower growth in *ex vivo* CSF compared with isolates derived from HIV-infected patients, with median fungal burdens of 7×10^3^ CFU/ml (IQR4.9 ×10^3^ - 1.0×10^4^) versus 1.9×10^4^ CFU/ml (IQR 6×10^3^ - 2.6×10^4^) after 48 hours, P=0.02 (Figure 2). The isolates derived from immunocompetent patients also elaborated significantly thinner capsules (median thickness 1.3 microns, IQR 0.8 - 1.6) compared with isolates from HIV-infected patients (median 1.5 microns, IQR 1.3 – 1.95), P<0.001).

### Low virulence VNIa-5 C. neoformans isolates can be induced to become highly virulent

We hypothesized that the development and/or maintenance of the increased virulence state was due to inter-yeast communication. To test this hypothesis we cultured an environmental VNIa-5 isolate of low virulence in media supplemented with sterilised culture filtrate (sCF) extracted from culture of VNIa-5 isolates with high virulence from immunocompetent patients. Virulence phenotypes were again compared in *Galleria*. Culture in media supplemented with sCF from all four organisms from immunocompetent patients resulted in increased virulence of the environmental isolates in comparison to their ‘naïve’ state; the hazard of death after induction increased between 2.2 and 2.9-fold (Figure 3). This change in virulence phenotype of the induced environmental isolates was stable over multiple *Galleria* infection cycles **(Figure S2)**. Conversely, culture in media supplemented with pooled sCF from HIV-associated VNIa-5 isolates had no significant effect on the virulence phenotype of environmental isolates (HR1.4, 95CI 0.8-2.3, P=0.2, ‘induced’ isolate versus naïve isolate; Figure 4).

**Figure 3:**
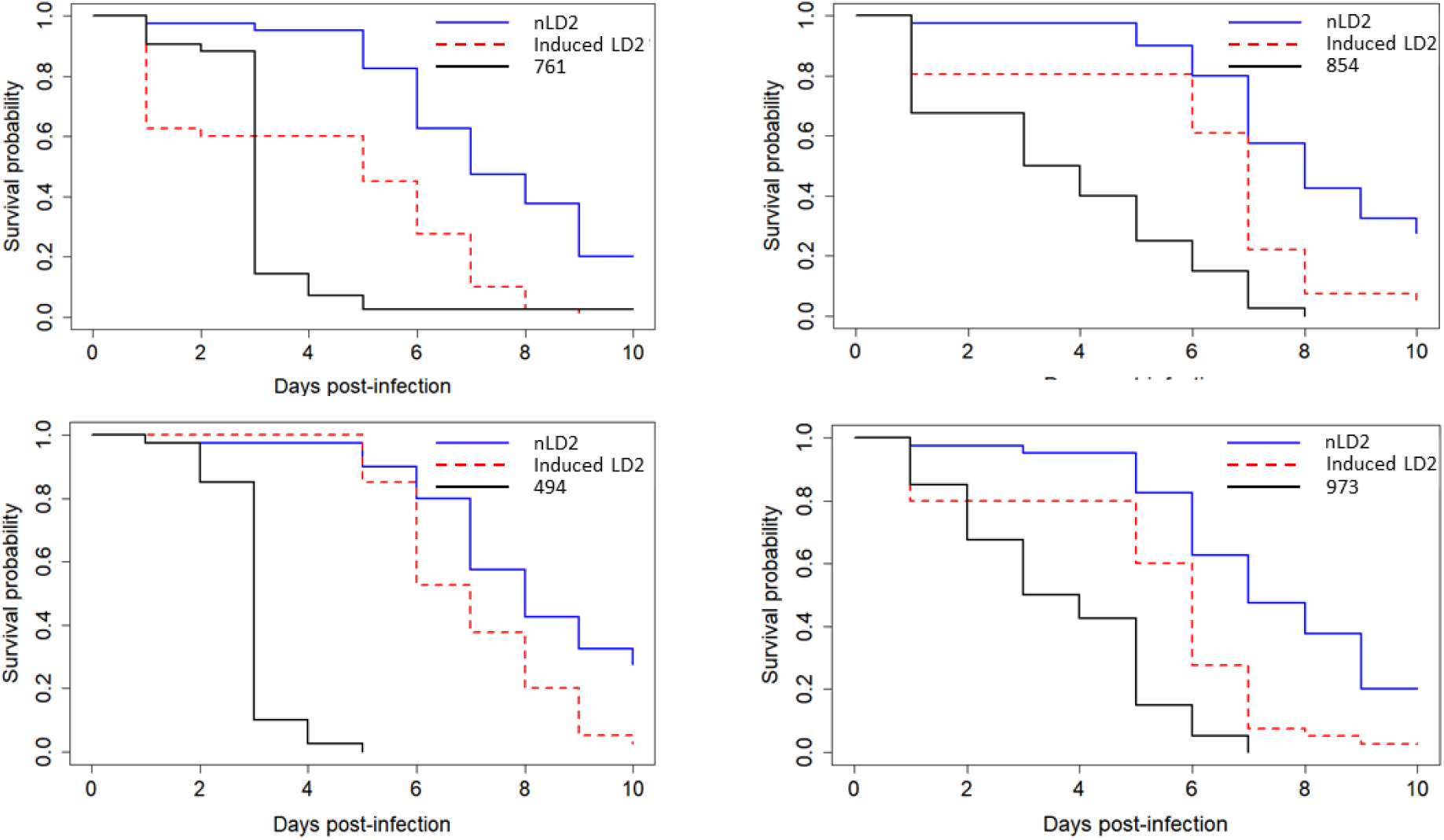
Survival curves for *G. mellonella* infected with 4 human isolates of *C. neoformans* lineage VNIa-5 derived from HIV-uninfected patients (BMD761, BMD854, BMD973 and BMD494), and naïve environmental strain (nLD2) and nLD2 following growth in sterile culture filtrate from each of the human derived isolates (induced LD2). Hazard ratios of the risk of death, 95%CI and P values for iLD2 versus nLD2 infections for each experiment are: HR 2.9, 95CI 1.8, 4.8 P<0.001; HR 2.5, 95CI 1.5, 4.0 P<0.001; HR 2.7, 95CI 1.7, 4.4 P<0.001 and HR 2.2, 95CI 1.4,3.6 P=0.001 respectively.

**Figure 4:**
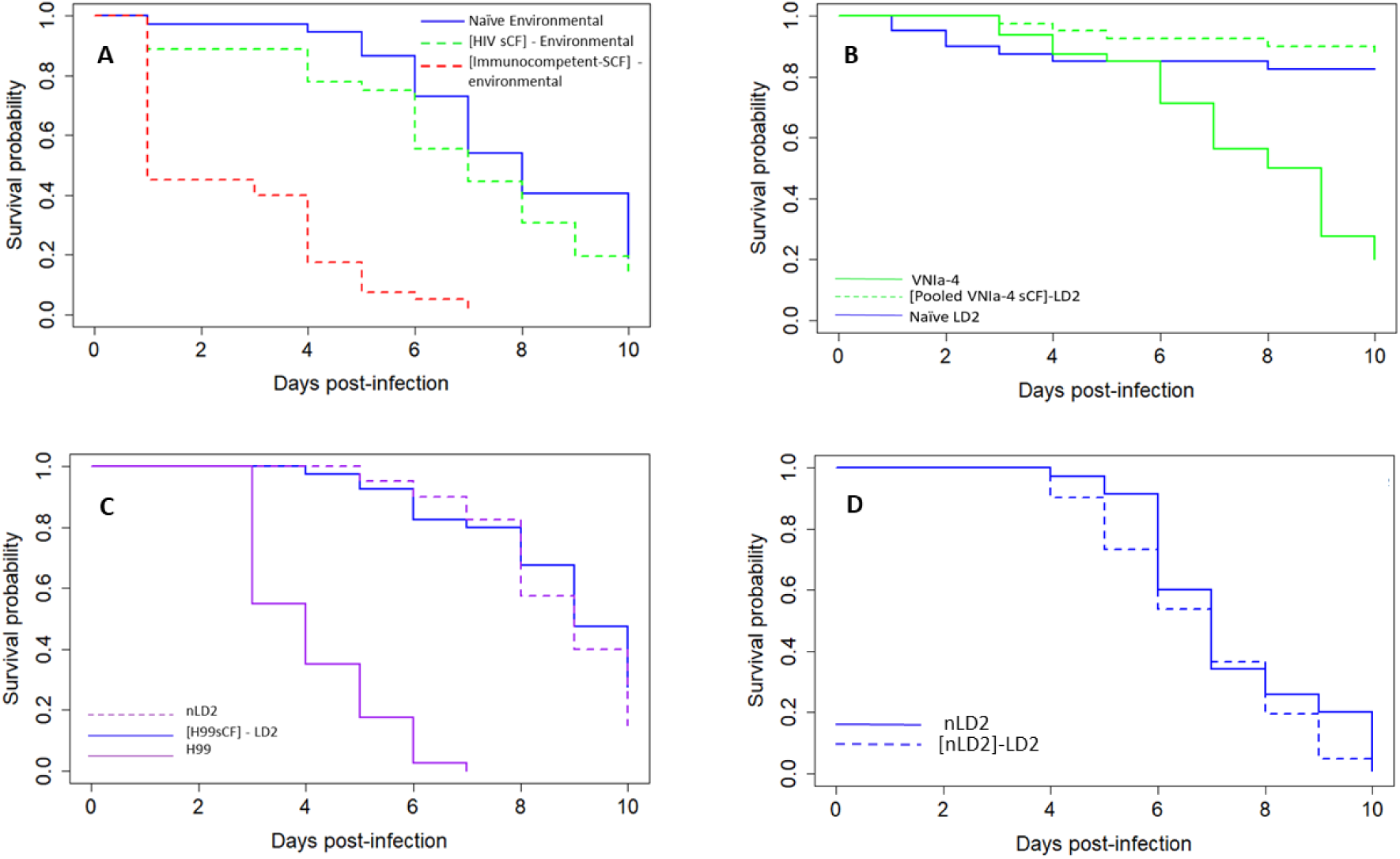
Survival curves of *Galleri*a infected with naïve environmental isolate, or that environmental isolate following growth in media supplemented with sterile culture filtrate (sCF) of different sources, or the source of sCF. The square brackets refer to the source of the sCF. **Figure 4A: Survival curves for *G. mellonella* infected with environmental isolates of *C. neoformans* lineage VNIa-5 (naïve environmental, blue line), and infected with the same environmental isolates following their growth in media supplemented with pooled sCF from either VNIa-5 lineage isolates derived from HIV-infected patients ([HIV-sCF], green dashed line) or from immunocompetent patients ([Immunocompetent-sCF], red dashed line)**. There is no significant increase in the hazard of death following infection with environmental isolates grown with sCF from HIV associated VNIa-5 isolates (HR 1.4 (95CI 0.8-2.3), P=0.2). Growth in media supplemented with sCF from isolates from immunocompetent patients is associated with a significant increase in the hazard of death (HR 10.0; 95CI 5.6-17.9) P< 0.001 sCF-immunocompetent vs naïve. **Figure 4B: Growth of naïve environmental strain nLD2 in media supplemented with sCF from HIV derived VNIa-4 lineage isolates does not result in increased virulence in the *Galleria* model**. The naïve LD2 strain was grown in media supplemented with pooled sCF from isolates BK80 or BK224 (VNIa-4 lineage strains derived from the cerebrospinal fluid of HIV-infected patients). Infection with the LD2 environmental isolate grown in sCF from isolates of the VNIa-4 lineage did not alter the hazard of death compared with infection with LD2 (HR 0.6; 95CI 0.2, 2.0, P=0.5. **Figure 4C: Growth of naïve environmental strain nLD2 in media supplemented with sCF from the *Cryptococcus neoformans* H99 type strain does not result in increased virulence in the *Galleria* model**. H99 was significantly more virulent than naïve DL2 (HR 28.7 (14-60.4) and P<0.001 for H99 vs LD2) but there was no change in the hazard of death for *Galleria* between infection with either the naïve or H99 induced environmental isolate HR 1.3 (0.8-2.2) and P=0.25 for iLD2 vs nLD2 N = 40 larvae per arm. **Figure 4D. The virulence of the LD2 environmental isolate is not increased by growth in media supplemented with sCF derived from previous culture of its naïve state** (HR 1.4, 95% CI 0.9-2.2, P=0.2).

We repeated the above experiment but cultured an environmental VNIa-5 isolate in media supplemented with sCF from VNIa-4 isolates (necessarily) derived from HIV-infected patients. These conditions did not result in any significant variation in virulence. Similarly, culturing the VNIa-5 environmental isolate in media supplemented with sCF from a previous culture of the same organism, or from the H99 type strain, had no effect on virulence (Figure 4).

### Culture filtrate from an induced VNIa-5 isolate will increase virulence in its naïve isogenic self

We next measured the effect of growing a ‘naïve’ low virulence VNIa-5 environmental isolate (LD2) in media supplemented with sCF from its previously ‘induced’ higher virulence self. We cultured the naïve LD2 isolate in culture medium supplemented with sCF from a highly virulent isolate derive from an immunocompetent patient (BMD761) for 48 hours. The cultured yeast cells were harvested by centrifugation, washed, and innoculated onto solid media for single colonies. We termed this isolate ‘iLD2’. iLD2 was again cultured in liquid media for 48 hours and the culture filtrate harvested. The naïve LD2 isolate was cultured in media supplemented with the sCF from iLD2 for 48 hours. We termed the resulting isolate ‘iLD2-induced LD2’. The virulence phenotypes of the isolates from each experiment in the *Galleria* model – i.e. ‘LD2’, ‘iLD2’, ‘iLD2-induced LD2’ and BMD761 were compared. This experiment demonstrated that, like sCF from BMD761, sCF from iLD2 was itself able to induce an increased state of virulence in its naïve self (nLD2): HR 2.8 (1.7-4.4), P<0.001 (Figure 5).

**Figure 5:**
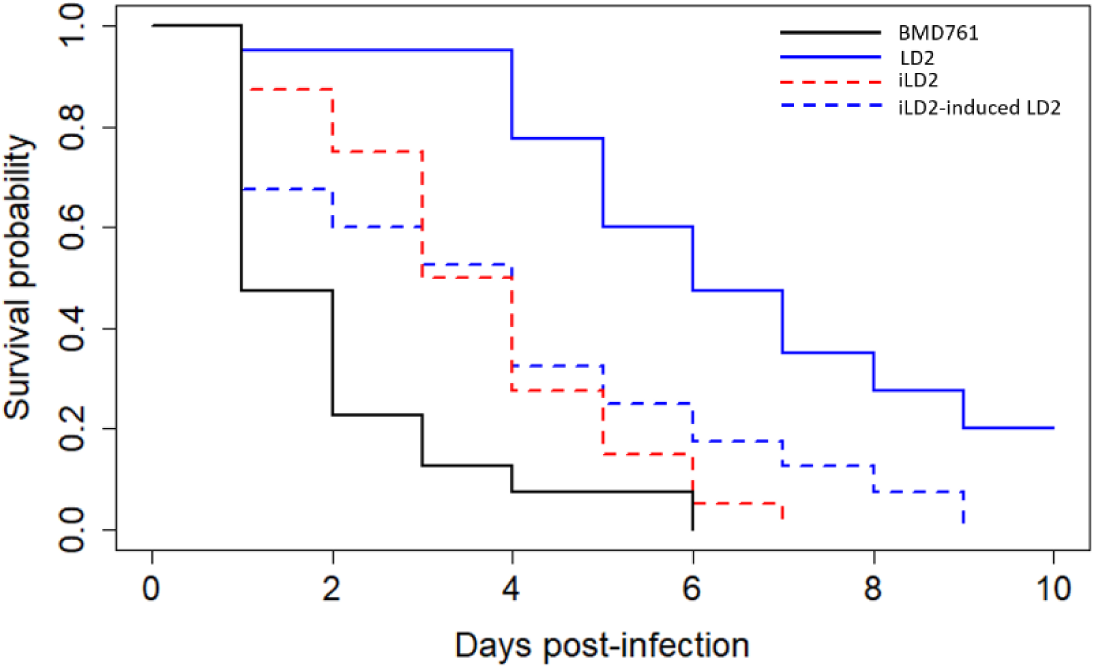
Survival curves for *Galleria* infected with one of 4 VNIa-5 isolates: LD2: ‘naïve’ environmental isolate; BMD761: isolate from immunocompetent patient (HR 8.2 (4.9-13.8), P<0.001 vs LD2); induced LD2: naïve LD2 isolate that has been grown with sCF (sCF) from BMD761 (HR 3.6 (2.2-6.0), P<0.001 vs LD2); iLD2-induced LD2: naïve LD2 isolate grown with sCF from iLD2 (HR 2.8 (1.7-4.4), P<0.001 vs LD2). N=40 larvae per arm

### Virulence induction is associated with known virulence markers

We assessed the effect of culture filtrate induction on known virulence-associated phenotypes in *Cryptococcus* by comparing the environmental isolated LD2 before (naïve) and after induction (induced). There was no change in thermotolerance or melanization. However, the induced LD2 had superior growth after 48 hours incubation in *ex vivo* CSF compared with its naïve self (median 5.9 ×10^4^ CFU/ml, IQR 3.6×10^4^ – 7.6×10^4^ vs 3.5 ×10^4^ CFU/ml, IQR 2.3 – 5.7 ×10^4^, P= 0.02, Figure S3a). Similarly, the 48 hour fungal burden in hemolymph of *Galleria* infected with the induced isolate was significantly higher than in hemolymph when infected with its naïve self (median fungal burden 9.6 ×10^6^ CFU/g body weight (inter-quartile range 6.9 ×10^6^ - 1.4×10^7^) versus 5.3 ×10^6^ CFU/g body weight (IQR 4.0 ×10^6^ - 6.7 ×10^6^, P=0.025), Figure S3b). Induced LD2 isolates eloborated significantly thinner capsules compared with the naïve self both when grown *in vitro* (median capsule width 4.5 µm, IQR 3.3-6.4 versus 7.2 µm, IQR 4.5 – 9.8, P<0.001, induced versus naïve respectively) and when recovered from larval hemolymph (P=0.03, see Figure S4). Both induced LD2 and naïve LD2 had significantly thinner capsules than the highly virulent immunocompetent associated organism BMD761 (P<0.001 and P=0.03 respectively).

### The induction phenomenon is protein mediated via extracellular vesicles

The induction effect of sCF from hypervirulent BMD761 on the naïve environmental organism LD2 was maintained after storage at -20°C for 28 days. The same observation was made for sCF treated with RNase or DNase. However, boiling sCF, or treating sCF with Proteinase K, abolished the induction effect (Figure 6). We further hypothesized that the induction phenomenon was associated with extracellular vesicles (EVs), which have been implicated in the pathogenic potential of *Cryptococcus neoformans* and the related organism *Cryptococcus gattii*^13-15^. Consequently, we added bovine serum albumin (BSA), which disrupts EV function, to sCF from BMD761 and measured the induction effect on LD2^13,16^. Adding BSA resulted in the loss of the induction effect with no increase in the virulence of the recipient isolate versus its naïve self (HR 0.8; 95CI 0.5, 1.3, P=0.32, Figure S5).

**Figure 6:**
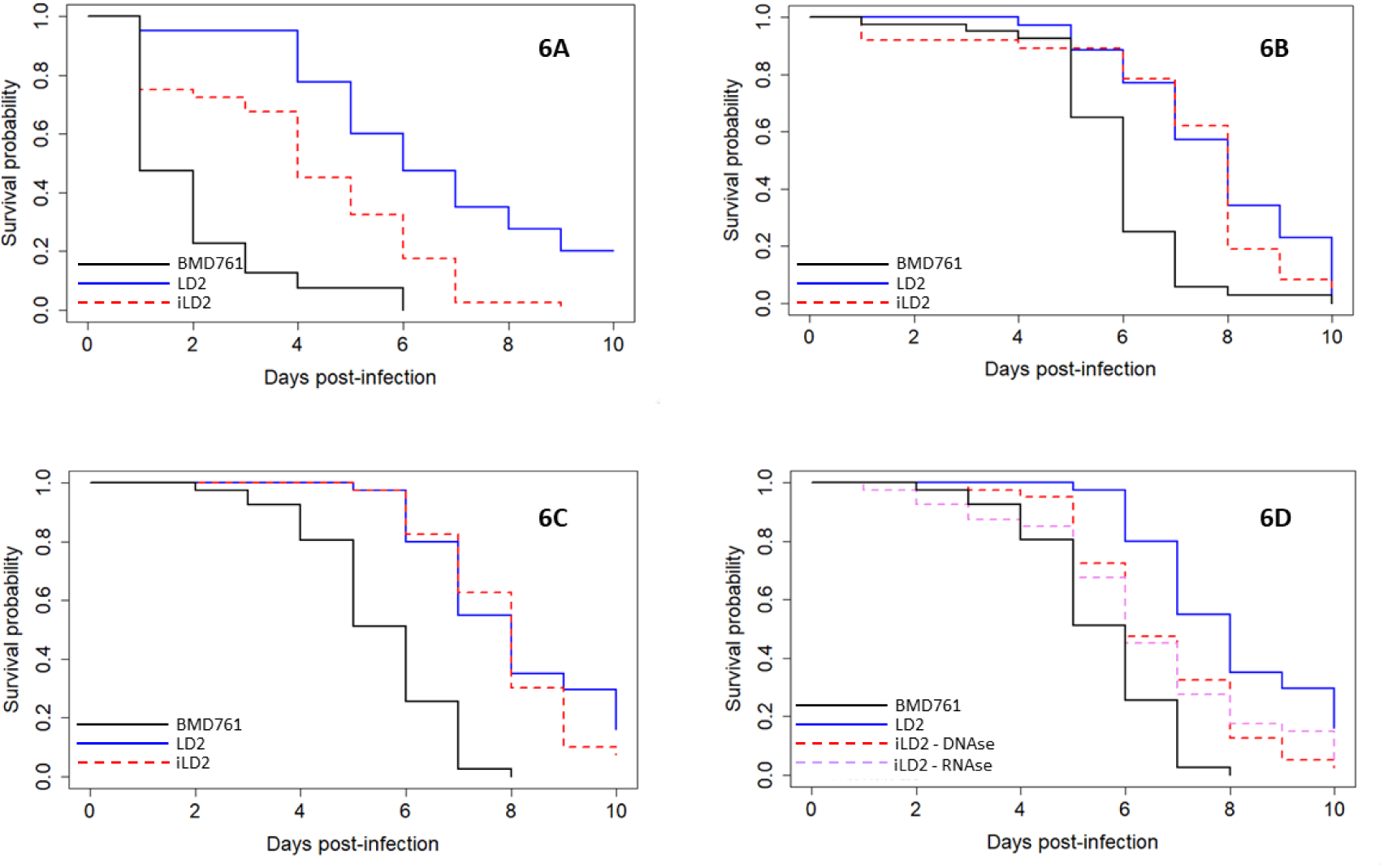
The effects of treating sterile culture filtrate from highly virulent strain BMD761 with either freezing, boiling, protease or nuclease on its ability to induce increased virulence in the naïve environmental strain LD2. Each figure shows survival curves for *Galleria* infected with different *C. neoformans* isolates. BMD761 is the high virulence VNIa-5 lineage isolate derived from an HIV-uninfected patient, LD2 is a low virulence VNIa-5 lineage isolate derived from the environment, and induced LD2 is the naïve environmental LD2 isolate following growth in media supplemented with sCF from BMD761. **Figure 6A: The inducing effect of sCF is not affected by freezing at -20C**. HR 2.7 95%CI 1.7, 4.5 P<0.001 **Figure 6B. Boiling abolishes the induction effect of sCF**. HR1.3 95%CI 0.8, 2.01, P=0.3. **Figure 6C**: **The inducing effect of sCF is abolished by treatment with proteinase**. HR1.2, 95CI 0.8, 2.0, P=0.36 **Figure 6D. Treatment with DNAse or RNase has no effect on the induction effect of sCF**. DNAse HR 2.1, 95CI 1.3, 3.4, P=0.002; RNAse HR 2.0, 95CI 1.2, 3.4, P=0.004. N = 30 *Galleria* per arm for all experiments.

We then generated EV-free supernatant and EV-containing pellet fractions (as shown by electron microscopy) from both BMD761 derived sCF and previously induced LD2 sCF (Figure 8). Incubation of naïve LD2 in media supplemented with supernatants did not result in induction of increased virulence (BMD 761 supernatant: HR 1.2, 95CI 0.8, 1.5; P=0.3, Figure 7B and iLD2 supernatant: HR 1.2, 95CI 0.9, 1.6; P=0.1 see Figure 7D). Conversely, when a naïve LD2 was grown in media supplemented with the EV-containing pellets there was induction of increased virulence (BMD 761 pellet: HR 1.8, 95CI 1.4, 2.4; P<0.001, Figure 7A and iLD2 pellet: HR 2.1, 95CI 1.6, 2.7; P<0.001, see Figure 7C).

**Figure 7:**
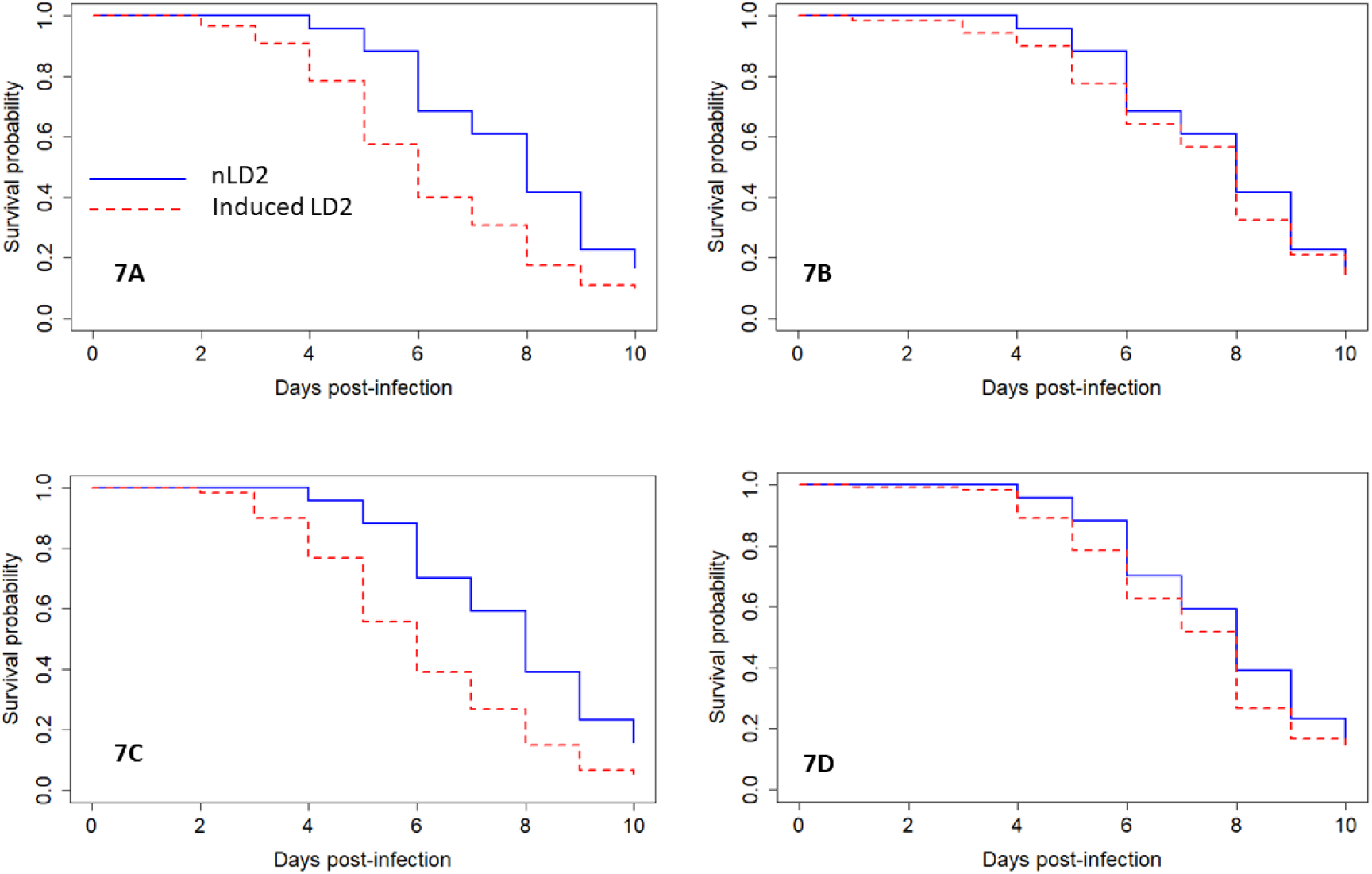
The effect of ultracentrifugation of sterile culture filtrate on induction of virulence in naïve environmental isolate nLD2. SCF, derived either from highly virulent isolate BMD761, or from LD2 strain following virulence induction with sCF from BMD761, underwent an extracellular vesicle (EV) isolation protocol to produce a putatively EV containing pellet, and an EV-free supernatant. These fractions were then added to broth culture of naïve LD2 isolate, and the relative virulence of naïve LD2 was compared with subsequent ‘induced LD2’ (iLD2). Experiments were done in triplicate, N=80 *Galleria* per arm. Summated data shown. **Figure 7A: Growth of LD2 in media supplemented with ultracentrifugation pellet derived from BMD761 sCF**. Induction of increased virulence is seen, HR 1.8, 95CI 1.4, 2.4; P<0.001. **Figure 7B: Growth of LD2 in media supplemented with ultracentrifugation supernatant derived from BMD761 sCF**. No induction of virulence in LD2 is seen, HR 1.2, 95CI 0.8, 1.5; P=0.3. **Figure 7C: Growth of LD2 in media supplemented with ultracentrifugation pellet derived from sCF from LD2 that has previously been induced with BMD761 sCF**. Induction of increased virulence is seen, HR 2.1, 95CI 1.6, 2.7; P<0.001. **Figure 7D: Growth of LD2 in media supplemented with ultracentrifugation supernatant derived from sCF from LD2 that has previously been induced with BMD761**. No induction of increased virulence is seen, HR 1.2, 95CI 0.9, 1.6; P=0.1. In all panels the solid blue line represents *Galleria* infected with naïve LD2 (nLD2) and the red dashed line represents *Galleria* infected with the ‘induced’ LD2 (iLD2), i.e. the environmental isolate following growth in media supplemented with either pellet or supernatant.

**Figure 8:**
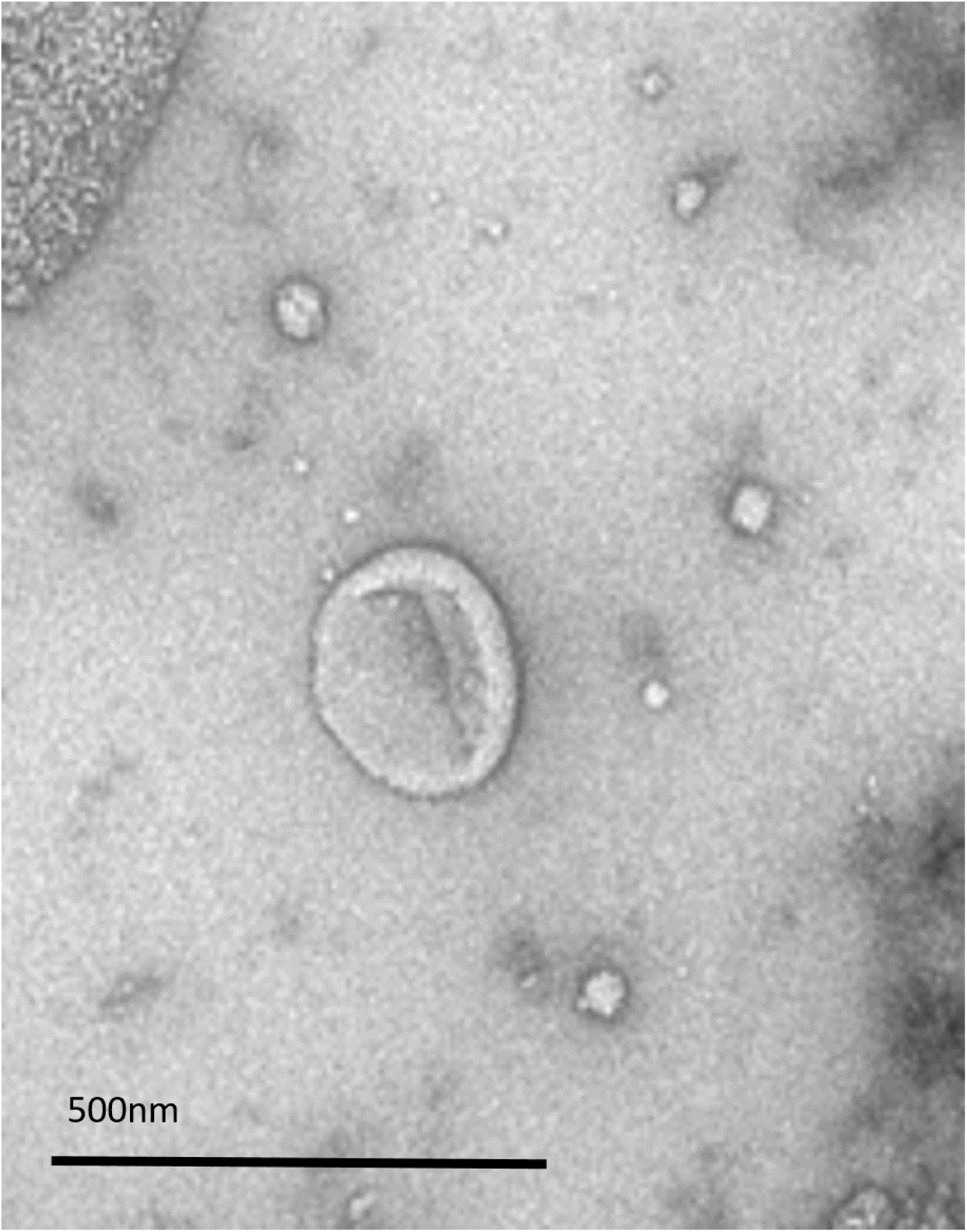
Transmission electron micrograph showing example of extracellular vesicle in ultracentrifugation pellet of BMD761.

### Induction is associated with the expression of genes encoding virulence factors and hypothetical proteins

We used RNA-seq to investigate the transcriptional basis of the observed phenotypic change. High quality RNA sequences were generated from four organisms with a confirmed phenotype: an environmental VNIa-5 (low virulence and termed naïve LD2 (nLD2)), a clinical VNIa-5 from an HIV-uninfected patient (BMD761), an environmental VNIa-5 in a state of increased virulence after growth in media supplemented with sCF from clinical VNIa-5 (induced LD2: iLD2) and the naïve environmental VNIa-5 isolate (nLD2) grown in media supplemented with its own sCF from a previous culture of its naïve self (self-induced LD2: siLD2) (Table S2).

The majority of transcriptome replicates clustered by their sample type on a PCA plot of read counts per gene (Figure S6). There was a notable difference in global gene expression between BMD761 and nLD2. The overall transcriptional profile of iLD2 was closer to that of BMD761 after induction. The ‘self-induced’ LD2 also differed in the transcriptional profile from the naïve LD2, was distinct from iLD2, and did not converge with the expression profile of BMD761.

We next performed differential gene expression analysis to identify genes that were significantly (Wald test, Benjamini-Hochberg adjusted P value <0.05, log2 fold change ≥1 and ≤-1) up or down regulated. We filtered 784 differentially expressed genes between BMD761 and naïve LD2; after induction with BMD761 sCF; 244 genes were differentially expressed between BMD761 and induced-LD2 (Figure 9). This contrasts with the large number of differences between these 2 isolates and LD2 in the other states (naïve or self-induced), where we found there were >1500 DEGs (Figure 9).

**Figure 9:**
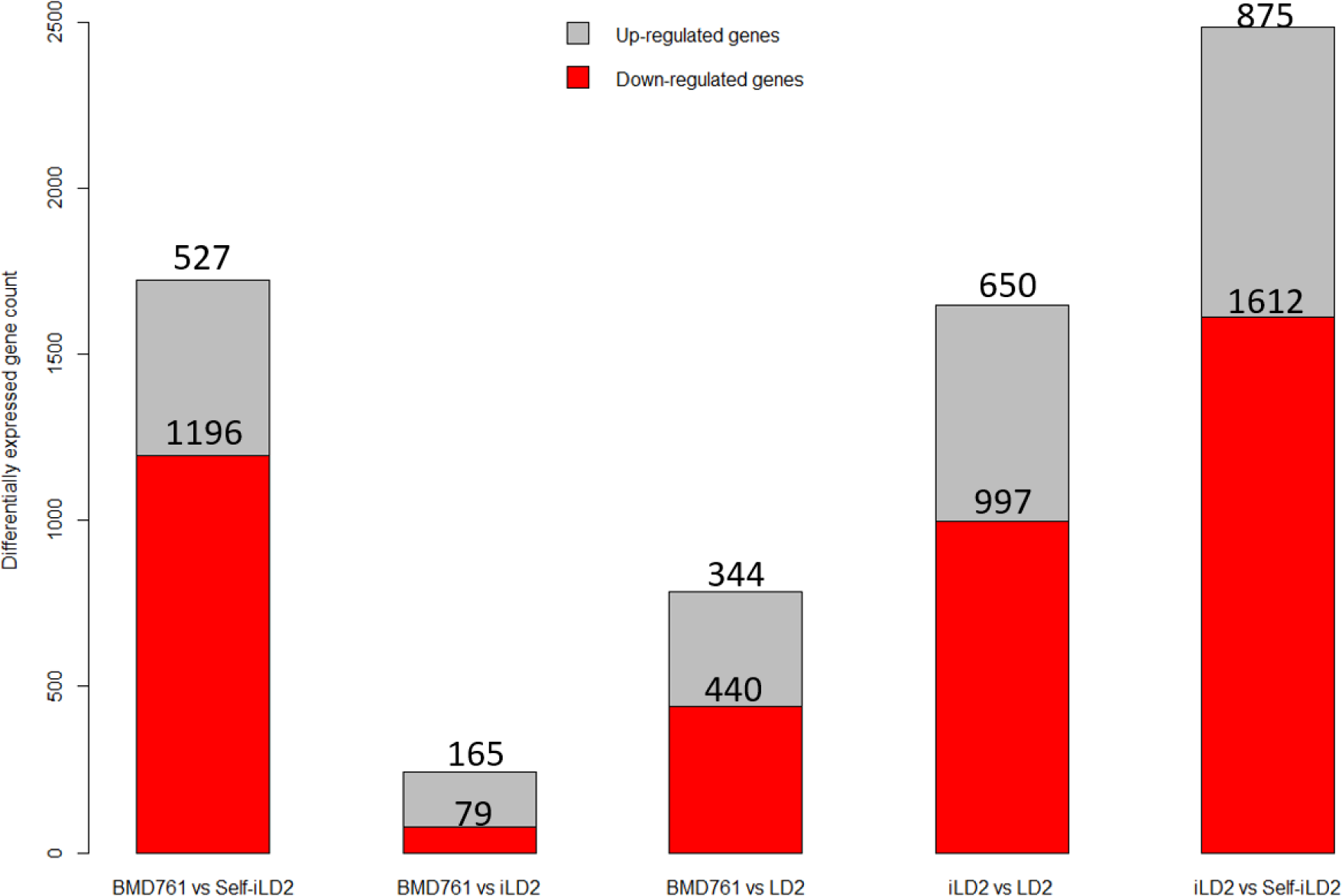
The number of differentially expressed genes between different samples (outliers removed). Genes were counted as differentially expressed if they had a Benjamani-Hochberg adjusted P-value of <0.05, and log2 fold change ≥1 and ≤-1.

We aimed to identify upregulated genes in both BMD761 and induced LD2 compared with naïve LD2 that may explain the phenotypic difference. Focussing on genes which were experimentally known to be associated with virulence^17^, we identified 17 up-regulated virulence genes in both BMD761 and induced LD2 relative to naïve LD2 (Supplementary Table S3). Specfically, we found that QSP1, which encodes a secreted peptide that has previously been implicated in quorum sensing and the upregulation of virulence of *C. neoformans*, was significantly downregulated in both the highly virulent BMD761 isolate and the induced environmental isolate versus its naïve self (Supplementary Table S3)^18^.

## Discussion

A number of lineages of *Cryptococcus neoformans* cause meningitis in humans in Vietnam, but the vast majority of cases are due to either VNIa-4 or VNIa-5^2^. We have previously shown that the VNIa-5 lineage is strongly associated with disease in apparently immunocompetent patients^2,3^. Both VNIa-5 and VNIa-4 frequently affect our (severely immunosuppressed) HIV-infected patients^2,3^. VNIa-4 rarely causes disease in HIV-uninfected patients, and when it does, the patients are significantly more likely to have underlying diseases associated with immunosuppression^9^. Despite this difference in ability to infect hosts of different immune competencies, mortality rates are no different in HIV-infected patients whether they are infected with VNIa-4 or VNIa-5^3^. Our work here goes some way towards explaining these observations. We have found that the VNIa-5 lineage displays significant variability in virulence phenotype in the *Galleria* model of infection, and that this is associated with the particular isolate’s ecological background – isolates from immunocompetent patients are more virulent than those from HIV-infected individuals or those recovered from the environment. The VNIa-5 isolates derived from immunocompetent patients are also significantly more virulent than VNIa-4 isolates, necessarily derived from HIV-infected patients.

The differences in virulence phenotype of the VNIa-5 isolates are not explained by a specific within-lineage substructure - virulent isolates from immunocompetent patients do not cluster within, but are distributed throughout, the VNIa-5 phylogeny^2^. This suggests that any isolate of the VNIa-5 lineage should have the capacity to express a highly virulent phenotype, and therefore infect immunocompetent patients. Consistent with this, we found that the VNIa-5 virulence phenotype is highly plastic – less virulent isolates, derived from the environment, can be induced into a more virulent state by growing them in culture media that has been supplemented with sCF from highly virulent isolates, derived from immunocompetent patients. While we found HIV derived VNIa-5 isolates were somewhat more virulent that the environment-derived isolates, their sCF lacked this capacity and could not induce increased virulence. The induction effect was also limited to sCF derived from the VNIa-5 lineage – sCF from the VNIa-4 lineage, or from the VNIb H99 type strain, did not have the induction property. Therefore, we hypothesize that the induction effect seen with culture filtrate from immunocompetent patient-derived VNIa-5, and the virulence plasticity of the lineage, are intimately associated with the ability of the lineage to cause disease in the immunocompetent host.

We found the increased virulence of VNIa-5 derived from immunocompetent patients was associated with differences in *in vitro* virulence phenotypes compared with isolates from HIV-infected patients, although the relationship was not simple. Isolates from immunocompetent patients had faster rates of growth at 30 and 37°C, but had slower growth in *ex vivo* cerebrospinal fluid and expressed thinner capsules. Induction of the higher virulence state in the environmental isolate LD2 was also associated with increased fungal burdens in *Galleria*, and expression of thinner capsules both *in vitro* and in the *Galleria* model. However, the induced LD2 isolate grew more rapidly in *ex vivo* CSF compared with its naïve self. It is not clear why the changes in *in vitro* phenotypes on induction were not entirely consistent with our findings when comparing isolates by source. It may reflect the fact that these measures of virulence are largely validated in relation to the ability to cause disease in humans irrespective of immune background, i.e. including those with HIV infection. Or, it may be that the induction phenomenon we have identified here is only a part of the mechanism that is associated with disease in immunocompetent humans. However, our transcriptional data suggest thet the change in virulence we are measuring in *Galleria* is real and relevant, because i) it is associated with increased expression of a number of virulence associated genes and ii) we saw a significant and remarkable convergence in gene expression between the highly virulent BMD761 isolate and the induced environmental isolate LD2.

The induction phenomenon we see is robust and repeatable. We do not believe it is an artefact of ‘carry-over’ of virulent organism from the sCF donor isolate, because we used 0.45 micron filters, which should exclude any donor cells, and every culture filtrate was tested for sterility by culture. Furthermore, the induced state is stable following purification by culture on solid media. We can be certain that the convergence of the transcriptomic data from BMD761 and iLD2 is not confounded by contamination because we identified SNPs unique to each isolate within the context of nearly 900 *C. neoformans* genomes^2^, and verified the identity of the strains in the RNA-seq data using these markers. Similarly we also do not believe that the effect on virulence in *Galleria* is due to ‘carryover’ of some agent (eg extra-cellular vesicles) from the donor isolate sCF into *Galleria*, because induced isolates retain their increased virulence following purification on solid media.

The induction effect appears to include a protein or small peptide component, since the inducing effect of sCF is lost with both boiling and protease treatment. A secreted peptide – QSP1 – has previously been shown to be involved in quorum-sensing and virulence regulation of *C. neoformans*^18^. Interestingly, we found that expression of this gene is down-regulated when the low virulence environmental strain is induced into the high virulence state and thus it is unlikely that this gene explains the phenomenon we observed.

Our data also suggest that the induction effect is associated with extra-cellular vesicles – the addition of albumin, which disrupts *Cryptococcus* vesicles, to sCF inhibited virulence induction^16^. We used an ultracentrifugation protocol^13^ to isolate EVs and showed that the induction effect was maintained in the EV containing pellet, and lost in the EV-free supernatant. We confirmed the presence and absence of EVs in the pellet and supernatant respectively with transmission electron microscopy. The populations and compositions of EVs secreted by eukaryotic cells are complex. In addition to proteins they can contain small nucleic acids^13^. While we found no effect of nuclease treatment on the property of sCF, it is possible that we may have missed an effect because any small nucleic acids may have been protected within vesicles.

A striking feature of the induction effect is the stability of the resulting phenotype, and the transmissibility of the phenomenon. One possible explanation is that the effect we are observing is an infection event by a mycovirus. To investigate this possibility we interrogated all unmapped reads from our whole genome and transcriptome sequence data using a metagenomic platform ((http://taxonomer.com, data not shown). We found no evidence of viral infection. A further possibility is that the effect is prion mediated – an EV-associated prion (Sup35p) has been implicated in cell-to-cell communication in *Saccharomyces cerevisiae*, although the effect is not well-conserved across species^19,20^. However, the fact that the effect of sCF we observed was heat labile suggests a prion-driven mechanism is unlikely.

On balance, we believe the phenomenon we have described is mediated through EVs associated with one or more peptides/proteins. Small nucleic acids may also be involved. An EV-dependent mechanism is plausible - EV production has been demonstrated for a number of fungal pathogens, including *Cryptococcus*^14,15,21-23^. A specific role in the pathogenicity of *C*.*neoformans* was first postulated in 2008^22^, and EVs have been shown to contain virulence-associated factors including capsular components and laccase^14^. In addition to proteins, EVs are also associated with small DNAs and RNAs which could influence virulence^13,23-25^. More recently, EVs have been shown to mediate the virulence of an outbreak strain of the related species *C*.*gattii*^13^. The EV-mediated effect was dose-dependent, manifested as increased growth, and required both EV-associated protein and RNA, providing strong evidence of EV uptake enabling inter-yeast communication^13^. The phenomenon differed from our observations in that it was seen in macrophage culture, and onward transmissibility of the phenotype was not reported^13^.

Where and when the virulence induction phenomenon occurs in relation to human disease is unclear. It may be a rare event that occurs somewhere in the environment, perhaps as a result of interaction with other micro-organisms, or a consequence of animal infection. If this is the case, we would on occasion expect to see such inducing, high virulence isolates in HIV-infected patients. To confirm this we would need to characterise large numbers of isolates from HIV-infected patients. Alternatively, induction might occur in humans themselves, although if so this is probably not driven by innate immunity, since the virulence phenotype was extremely stable in *Galleria*^26-29^.

A great challenge in treating HIV-uninfected patients with cryptococcal meningitis is knowing when it is is safe to stop antifungal therapy. Cure may take many months of treatment, and patients may even require life-long antifungal suppressive therapy^30^. This contrasts with HIV-infected patients, who have a modifiable immune pathology - once antiretroviral therapy has allowed immune recovery antifungal treatment can be stopped safely^30^. Inter-yeast signalling systems, associated specifically with changes in virulence, are potential therapeutic targets. Disrupting such systems might allow shortened treatment courses or even cure in immunocompetent patients. However, they are only relevant if active in human disease. Currently, it is extremely difficult to obtain sufficient volumes of clinical samples to reliably test hypotheses concerning pathogenesis (generated from laboratory observations) at the site of human disease^31^. However, the recent development of a novel cerebrospinal fluid filtration technology – neurapheresis – may revolutionise this. Neurapheresis, developed to treat hemorrhagic stroke, can efficiently remove large amounts of pathogen biomass from the cerebrospinal fluid of rabbits experimentally infected with *Cryptococcus*^32-34^. Such a system, itself a potential treatment for human cryptococcal disease, could allow confirmation that the phenomenon we describe here is active in human disease.

## Conclusion

The virulence phenotype of *Cryptococcus neoformans* VNIa-5 in the *Galleria* infection model is associated with the isolate source. Isolates from immunocompetent patients are the most virulent; those from the environment the least. The virulence phenotype is highly plastic and moderated by inter-yeast signalling through the secretion of proteins or peptides associated with extracellular vesicles. Induction of virulence is accompanied by transcriptional re-modelling and convergence of gene expression between the induced and inducer isolates. The induction phenomenon is VNIa-5 specific, and likely key in the lineage’s ability to cause disease in the immunocompetent host. Better understanding this mechanism may reveal novel therapeutic targets.

## Funding

Funded by a Wellcome Trust Intermediate Fellowship to JND Grant number: WT097147MA

### Acknowledgements

Electron microscopy was performed at the Dunn School EM Facility at the University of Oxford.

## Methods

### Isolates

Clinical isolates were derived from Vietnamese patients with cryptococcal meningitis who had been enrolled into descriptive and randomised intervention studies performed by our institute in collaboration with the Hospital for Tropical Diseases, Ho Chi Minh City^9,35,36^. All isolates were from cerebrospinal fluid obtained at the point of diagnosis and/or study entry, and stored on Microbank beads at -80°C prior to revival on Sabouraud dextrose agar. Informed consent was obtained from all participants; all studies received ethical approval through the Oxford Tropical Ethics Committee, The Hospital for Tropical Diseases, and The Ministry of Health of Vietnam. All isolates have been previously sequenced and the data are publically available^2^. Isolates of the lineage and human immune phenotype of interest (HIV-infected or not) were randomly selected from the isolate database for subsequent experiments. Environmental isolates were obtained via randomised sampling of air, soil and tree bark in urban and rural areas within and around Ho Chi Minh City. All strains used were mating type alpha.

### *In vitro* phenotyping

#### Temperature – dependent growth

Growth at high temperature and in *ex vivo* human CSF were tested as previously described with modifications for quantitative assessment^37^. *Cryptococcus spp* were cultured on Sabouraud agar for 2 days at 30°C. Inocula were made from single colonies and adjusted to 10^8^ cells/ml, then serially 10-fold diluted and spotted in duplicate onto YPD agar in 5µl aliquots. Incubation was at 30°C or 37°C for 48 hours. After 48 hours, the number of colony forming units (CFU) was counted and expressed as CFU/ml.

#### Capsule production

Capusle thickness was assessed through growth on Dulbecco Modified Eagle Medium (DMEM) [supplemented with 4.5g/L glucose, L-glutamine, sodium pyruvate], NaHCO_3_ 250mM, NaMOPS 1M, Neomycin 200mg/ml, Cefotaxime 100 mg/ml^38^. Plates were incubated at 37°C in 5 % CO_2_ for 5 days. A suspension from a single colony was made in India ink visualized at 100X magnification using light microscopy (CX41, Olympus, Japan). Images were captured using a built-in DP71 camera (Olympus, Japan) and visualised using ImageJ (https://imagej.nih.gov/ij/index.html). Capsular thickness was calculated by subtracting yeast cell body diameter from the whole cell diameter. Ten individual microscopic yeast cells were assessed for each isolate.

#### Laccase activity

Laccase activity was assessed by adding an inoculum of 2×10^4^ cells into 96 microwell plates containing L-DOPA medium (0.1% glucose anhydrous, 0.1% L-asparagine, 0.3% KH2PO4, 0.025% MgSO4•7H2O, and 0.01% L-DOPA, pH 5.56) with incubation for 16 hours at 37°C followed by 48 hours at 25°C with shaking at 800 rpm to induce melanin production. After incubation, the supernatants were harvested by centrifugation (4,000 g for 5 minutes) and the amount of pigment produced determined spectrophotometrically at a wavelength of 475-nm. Absorabance was converted into laccase units (U) by a factor of 0.001 OD = 1 U^39^. A Two biological replicates were performed for each isolate.

#### Urease and phospholipase production

Urease production was confirmed by spotting 10μl of a 10^6^cells/mL cell suspension onto Christensen’s agar with incubation at 30°C. *Cryptococcus neoformans* H99 type strain and *Candida parapsilosis* ATCC 22019 were used as positive and negative controls. Cultures were observed daily for the characteristic pink discoloration.

Extracellular phospholipase activity was confirmed on egg yolk medium as previously described, with minor modifications again using a 5 µl aliquot of *C. neoformans* yeast suspension (10^6^ cells/ml) with incubation at 30°C for 72 hours^40^. The diameter of the precipitation zone (D) formed around a colony in relation to the diameter of the respective colony itself (d) was recorded for 5 selected colonies of each isolate (N=1450 total). The D/d ratio for each isolate was calculated. H99 was included for reference in each experimental batch. The final result for each isolate was expressed as the ratio D/d ratio. Type strain H99 was used as a quality control.

#### Survival in *ex vivo* cerebrospinal fluid

CSF was prepared by pooling and filtering (Millipore 0.45 micron membranes, Merck) CSF samples taken prior to antifungal therapy from HIV-infected patients participating in clinical trials. CSF was stored at -80°C prior to use. An inoculum of 10^6^ cells/mL of each isolate of interest was prepared using a Cellometer X2 (Nexcelom) cell counter. 10 uL of inoculum was added to 90 uL of CSF and PBS followed by incubation at 30°C for 3 days. After 72 hours the resulting culture was serially tenfold diluted, spotted onto Sabouraud plates and incubated at 30°C for 3 days for quantification. Tested trains were inoculated alongside H99 and the ΔENA1 negative control (which lacks a cation-ATPase-transporter resulting in decreased viability in human CSF and macrophages, strain provided by the Perfect Lab, Duke University)^37^. Two biological replicates were performed per isolate.

### *Galleria* infection model

Late instar isogenic wild type *Galleria mellonella* larvae (30 days from oviposition of the adult moth) were sourced from U U Animal, (Ho Chi Minh city, Vietnam). All larvae were kept at 16°C in bran in the dark. Larvae for experimentation were selected according to the following criteria: weight 250-300 mg, healthy colour (beige). Yeasts were revived from frozen stock (Microbank Beads) on SDA plates for 72 hours. Single colonies were picked and cultured in YPD broth for 24 to 48 hours in a shaking incubator (SI-300, Jeio Tech), at 150 rpm at 30°C until the concentration of cells was at least 10^8^ cells/mL. The pellet was collected with centrifuge at 8000 rpm in 1 minute and subsequently washed twice with PBS. The cell suspension was adjusted to a density of 10^8^ cells/mL using a Cellometer X2 (Nexcelom Bioscience, USA). Ten μL of inoculum (10^6^ cells/larva) were used for injection per larva. All inocula were relabeled by an independent person such that the operator/assessor was blind to the nature of the inoculum used in each batch of larvae (i.e. the survival experiments were blinded). Larvae were inoculated through injection using a sterile Hamilton syringe into the most caudal left pro-leg. 15-20 larvae were infected per cryptococcal isolate. Every survival experiment was internally contemporaneously controlled using larvae of the same batch, a negative control (PBS) (physical injury and sterility, data not show in survival plots), and uninjected larvae and the 2 or more experimental arms of interest. Infected larvae were incubated at 37°C for ten days and checked daily for mortality using physical stimulation with forceps.

### Culture Filtrate preparation

All isolates were grown in 7.5 ml YPD broth at 30°C with shaking for 48 hours followed by centrifugation at 8000 rpm for 1 minute. The resulting supernatant was filtered using a Millipore membrane filter 0.45 μm (Merck), and checked for sterility by plating onto Sabouraud’s agar and into BHI broth with incubation at 30°C for seven days.

### Induction experiments

2.5 mls of the culture filtrate of interest was added to 5mls of sterile YPD broth (total volume 7.5mls). A single purified colony of the isolate of interest was added to the resulting culture medium and incubated for 48 hours with shaking (150rpm) at 30°C. Cells were separated by centrifugation (1 minute at 8000 rpm), washed twice with PBS and pellet spread on Sabouraud agar plate for single colony purification. Single colony was then inoculated into YPD broth, and incubated with shaking for 48 hours for inoculum preparation.

### Confirmation of virulence phenotype stability in *Galleria*

Yeast was recovered from *Galleria* hemolymph, cultured for purity on SDA, recultured in YPD and reinoculated into *Galleria* as described 6 fold. Similarly purified culture derived from hemolymph was repeatedly subcultured on solid media, frozen on Microbank beads, revived, and re-inoculated into the *Galleria* model.

### Quantification of fungal burden in larvae

Larvae infected with the strains of interest (5 dead larvae/biological replicate) were collected at the point of death and gently squeezed to obtain 1 gram of hemolymph and fat body using a sterile knife. Glass beads (3mm) and 1 mL of sterile water were added for homogenization at 30Hz for 10 minutes (TissueLyser II, Qiagen). Homogenates were diluted with PBS and inoculated onto SAB plates and incubated for 3 days at 30°C. The number of colonies were counted to calculate the CFU/gram/larvae.

### Capsule size measurement from *Galleria*

Following infection with 10^6^ cells, larvae were sacrificed 48 hours post-infection and hemolymph extracted as above. Hemolymph was stained with India ink to visualise capsule using light microscopy (at 1000X magnification) with an Olympus DP71 digital camera (Olympus, Japan) and the proprietary Image-J software.

### Treatment of culture filtrate with freezing, heat, protease, and nuclease

To check the stability of culture filtrate exposed to prolonged frozen storage, the culture filtrate of BMD761 was kept at -20 °C for 33 days. The filtrate was brought to room temperature prior to induction experiments. To denature protein by heat, filtrate of BMD761 was boiled at 97 °C for 2 minutes. To denature protein chemically, filtrate was incubated with Proteinase K (Sigma-Aldrich, UK) at a concentration of 100 μg/mL in the presence of 30 mM Tris.HCl (pH=8) and 5 mM CaCl_2_ at 37 °C for 1.5 hours. To degrade DNA, 28 μL DNAse I (Ambion, ThermoFisher, UK) was added to 700 μL DNAse I buffer and 7 mL filtrate for 1 hour at 37°C. To remove RNA, 2.5 mL filtrate was incubated with 50 μL RNAse cocktail of RNAse A and RNAse T1 (Ambion, ThermoFisher Scientific, UK) for 1 hour at 37°C.

### Confirmation of digestion of protein and nucleic acid in filtrate

Protein digestion was confirmed by running treated and untreated culture filtrate on precast polyacrylamide gel 4-20 % (Mini-PROTEAN® TGX Stain-Free™ Precast Gels, BIO-RAD) at 200V for 40 mins. Precision Plus Protein Dual Xtra Standards (BIO-RAD) were loaded alongside samples, including a proteinase K treated ladder.

Samples treated with nuclease were confirmed by loading in agarose gel 1.5 % followed by electrophoresis for 60 mins at 100V, with 100bp DNA ladder.

### Extracellular vesicle preparation

Isolates for experimentation were grown to stationary phase in 500ml of YPD broth at 30°C with shaking for 48 hours prior to transfer to 200mL centrifugation vessels. These were spun at 15,000g for 10 minutes at 4°C. Supernatant was decanted and centrifugation repeated for a further 10 minutes, then filtered through 0.45um filter membranes. The filtrate was then concentrated using Amicon 100kDa ultrafiltration columns with centrifugation at 5000g for 15 minutes at 4°C. The EV containing retentate, was then ultracentrifuged at 150,000g at 4°C for 1 hour to provide EV containing pellet and EV-free supernatant.

### Electron Microscopy

Pellet and supernatant resulting from ultracentrifugation were fixed in 2% paraformaldehyde (PFA) in 0.1M sodium phosphate buffer at room temperature for 30 minutes. Samples were stored at 4 °C for up to 3 days before negative staining for Transmission Electron Microscopy (TEM). For negative staining, 10 µl of sample was applied to a freshly glow discharged carbon filmed 300 mesh Cu grid for 2 mins, blotted and stained with 2% uranyl acetate for 10 sec, then blotted and air dried. Grids were imaged in a Tecnai 12 TEM (Thermo Fisher) operated at 120 kV using a Gatan OneView camera.

### Preparation of inducing broth using ultracentrifugation product

Inducing broth consisted of 50uL ultracentrifuged product (either pellet or supernatant fraction) added to fresh sterile YPD to achieve a final volume of 7.5 mL. A 100uL volume was taken for culture as a test of sterility. Naive strains (a single colony each) were then inoculated into inducing broth and incubated for 48 hours with shaking at 30°C. Yeast were harvested in the normal manner via centrifugation 8000 rpm and washed twice with PBS prior to subculture on solid agar. Inoculum for larva infection (10^8^ cells/mL) were prepared in PBS using a Nexelom cell counter; 10 uL (10^6^ cells) were injected into the larvae. Inoculum densities were checked through serial dilution and culture.

### RNAseq

Three replicates of each strain were cultured in YPD broth for 96 hours at 30°C to determine log phase for RNA extraction. An inoculum of 10^6^ cells was grown in 7.5 mL of YPD broth by agitating (200 rpm). At interval time point post-inoculation (0, 6, 20, 24, 30, 44, 54, 68, 72 and 96 hours), an aliquot of 50 uL was taken for quantification by plating in Sabouraud agar plates. These plates were exposed to incubation at 30°C for 48 hours. Growth curves were plotted using ggplot package and R version 3.4.0 software (R Foundation for Statistical Computing, Vienna, Austria). Growth rates were determined using the formula growth rate=(log10N_t_-log10N_o_)*2.303/t-t_o_, where N indicates cell concentration at a particular time point; and t indicates the time point).

For transcriptional experiments, single colonies of each isolate of interest were revived from beads (Microbank, UK) stored at -80°C. Revived isolates (approximately 10^6^ cells, counted using a Cellometer) were grown in 7.5 ml of YPD broth by agitating for 18 hours at 30°C to reach mid-log phase. RNA was harvested using the RiboPure™-Yeast RNA Isolation Kit (Ambion, USA) according to the manufacturer’s instructions. Extracted RNA samples were subjected to quality control using an Agilent 2100 Bioanalyzer (Agilent, USA). The integrity of total RNA was estimated using the RIN (RNA Integrity Number), which ranges from 1 to 10, with 1 being the most degraded. Samples with RIN greater than 7 qualified for RNA-Seq. 6 biological replicates were prepared of every experimental condition. RNA was eluted in isopropanol and shipped at room temperature to Macrogen (Seoul, Korea) for library preparation and sequencing.

Paired-end RNA-Seq libraries were constructed using the TruSeq stranded mRNA preparation kit (Illumina, USA). Resulting cDNAs were ligated with sequencing adaptors and sequenced using the Illumina HiSeq 2000 platform, generating ∼11.2 million reads of 150 bp per sample, resulting in an estimated 177-fold coverage.

### RNA sequencing analysis

Raw reads were checked for quality using FastQC (https://www.bioinformatics.babraham.ac.uk/projects/fastqc/). Adapters were removed using scythe (https://github.com/vsbuffalo/scythe). The resulting reads were then further checked based on Phred score and read length in order to trim low quality regions using Trimmomatic (http://www.usadellab.org/cms/?page=trimmomatic). Sequenced reads were aligned with the *C. neoformans* var. *grubii* H99 reference genome using HISAT2 (https://ccb.jhu.edu/software/hisat2/index.shtml). SAM (Sequences Alignment Map) files, output of read alignment, were converted to BAM (Binary Alignment Map) files using SAMtools (http://www.htslib.org). Finally featureCounts was used to count reads assigned to genomic features (genes, exons, chromosome locations) on the reference genome^41^. The output table of raw counts per gene for each sample was imported for gene expression analysis using DESeq^42^. Library size differences were normalized internally^43^.

### Metagenomic analysis

Taxonomer, a metagenomics-based pathogen detection tool (available at http://taxonomer.com), was used to profile unmapped mRNA expression reads and unmapped DNA from (previous) genome sequencing from the *Cryptococcus* isolates. The web-based interface was used to visualize the summary of taxonomic composition and read abundance.

### Statistical data analysis

All statistical analyses were done using R software R version 3.4.2^44^. Kaplan-Meyer survival curves were plotted using ggplot2 and survminer package^45,46^. The Log-rank test was used to compare differences in survival. Hazard Ratios were estimated using the Cox proportional-hazards model. The Wilcoxon rank sum test was used to compare fungal burdens between isolates. Capsule size was compared using the Kruskal-Wallis test with P values adjusted for multiple comparisons with the Benjamini-Hochberg method.

## Supporting Information

### Supplementary Figures

**Figure S1:**
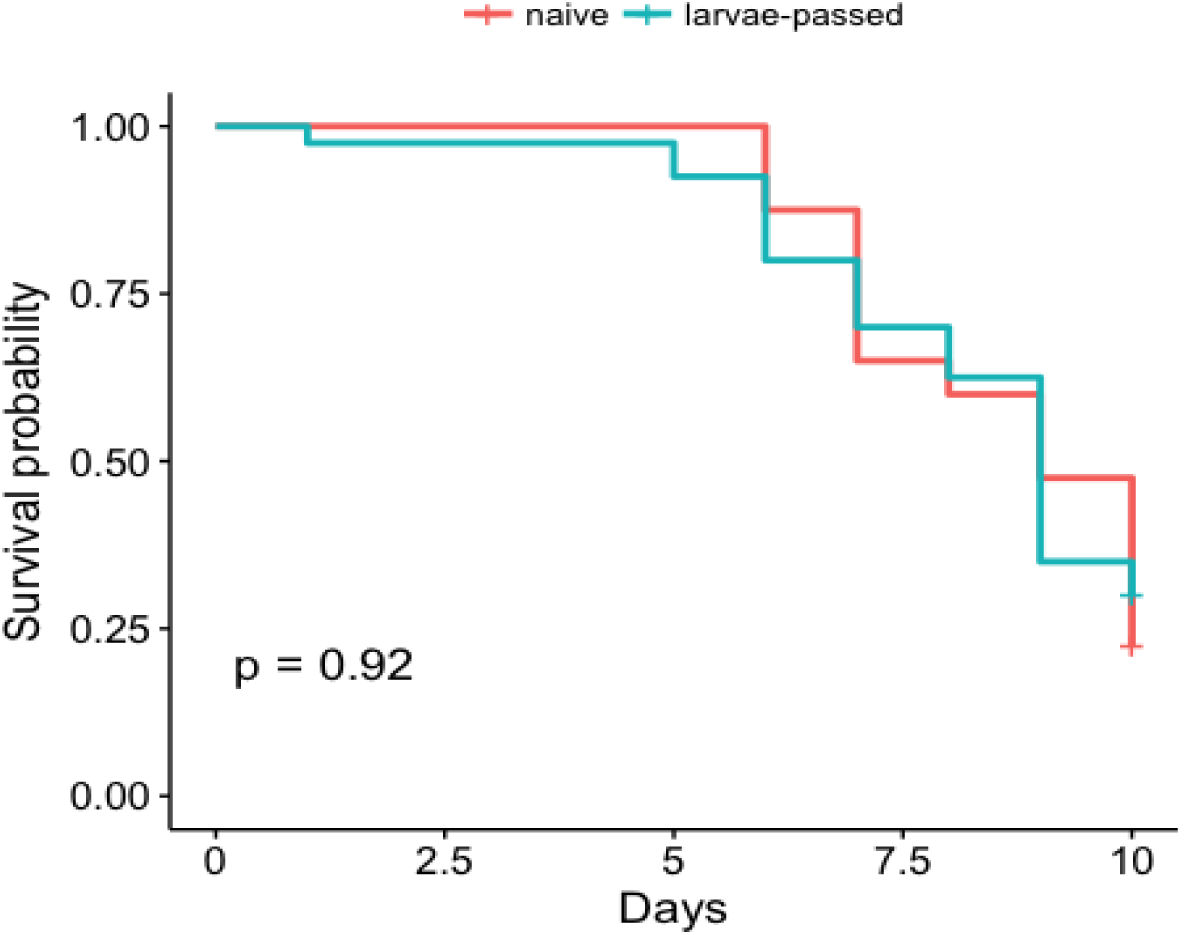
The virulence phenotype of environmental VNIa-5 isolate is stable over multiple infection passages (6-fold) through *Galleria* (P = 0.92, log rank test)

**Supplementary Figure S2:**
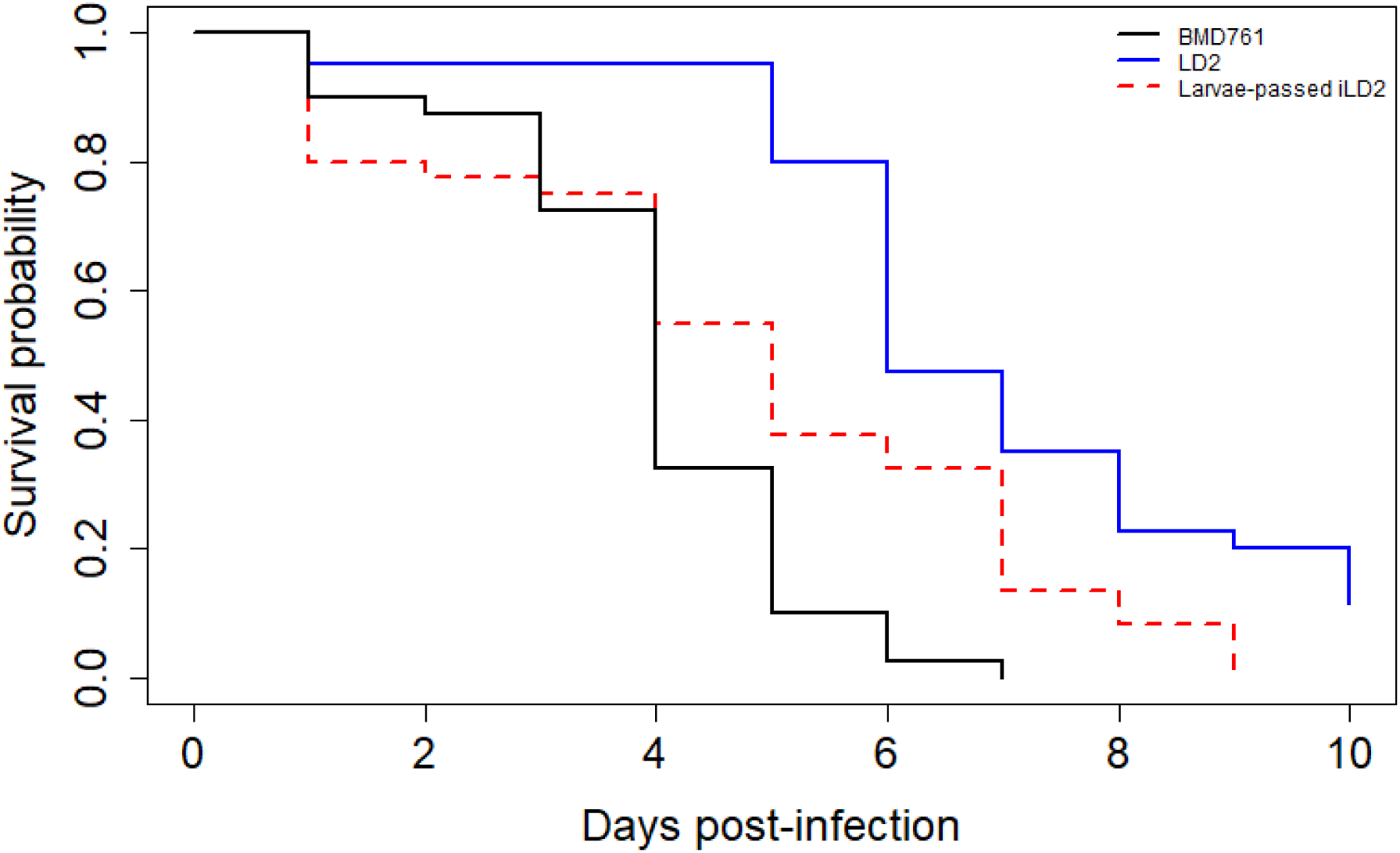
The induced increased virulence phenotype is stable over multiple generations. **Figure S2 shows survival curves for Galleria infected with one of 3 isolates:** The black curve is for Galleria infected with the hypervirulent immunocompetent patient derived BMD761 isolates, the blue curve for the naïve LD2 environmental isolate, and the red curve for infection with previously induced LD2 that has been repeatedly passaged through Galleria following recovery from haemolymph. The induced LD2 retains increased virulence compared with its naïve self. HR induced LD2 versus naïve LD2: HR2.3; 95CI 1.4, 3.7, P=0.001.

**Supplementary Figure S3:**
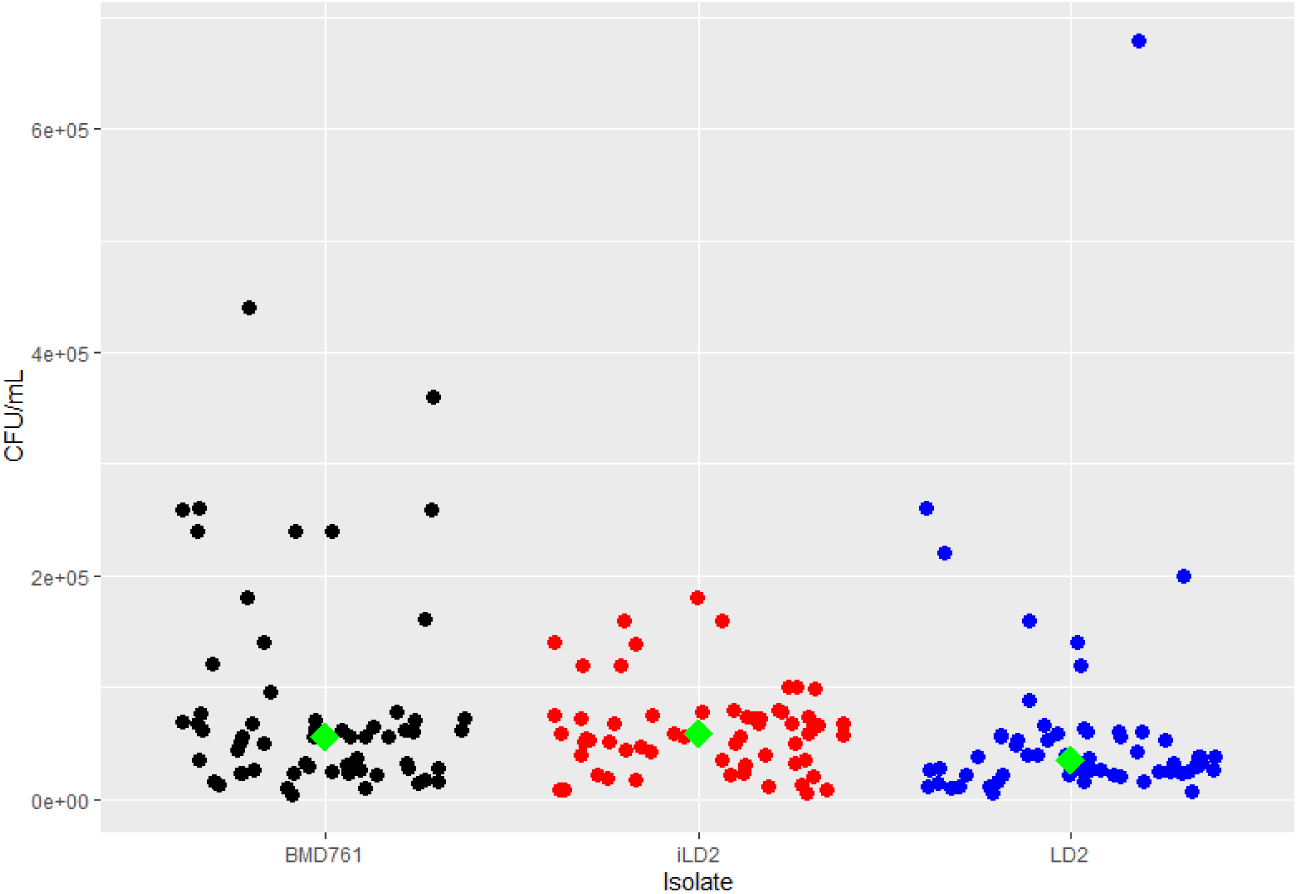
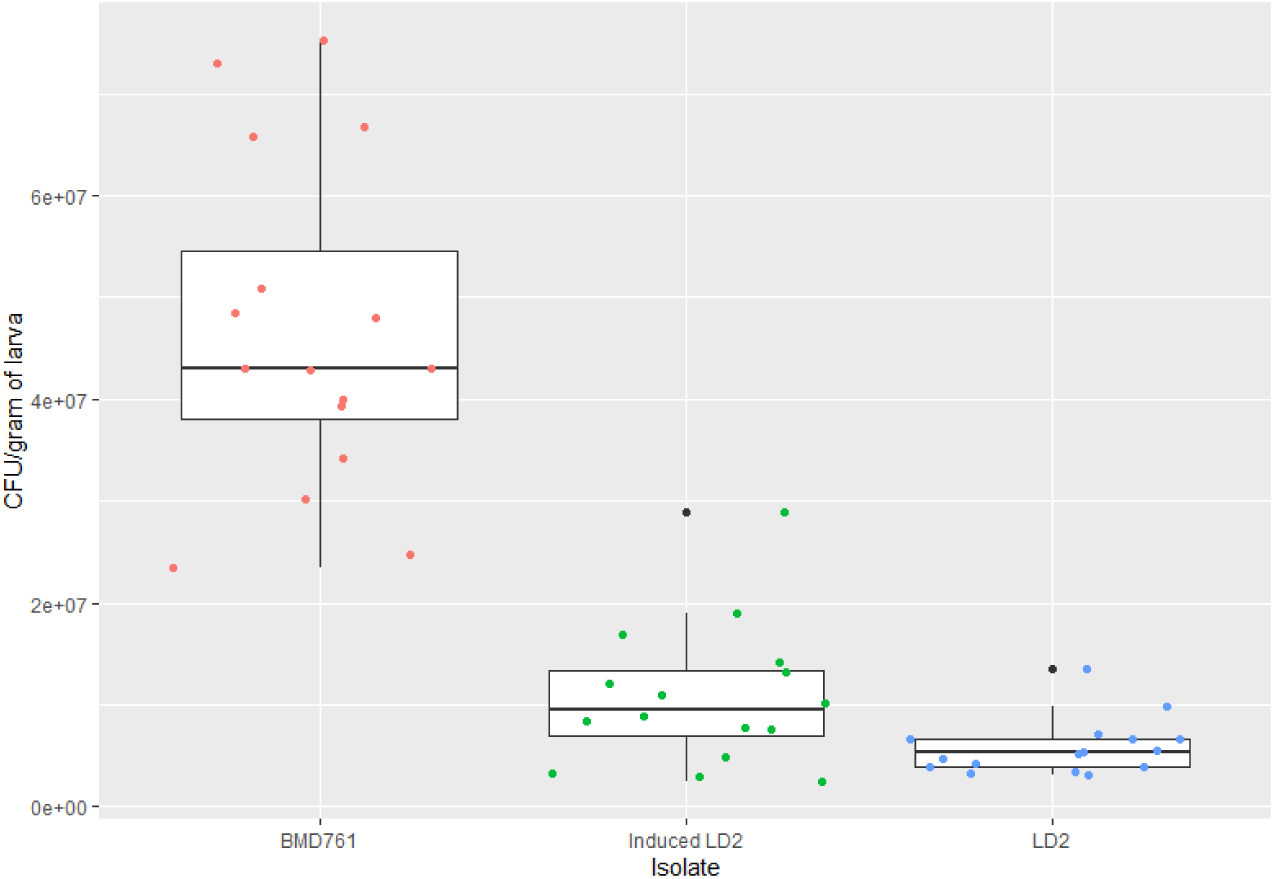
changes in *in vitro* virulence phenotypes following induction of nLD2. **a. Figure S3a: fungal burden ex vivo CSF 761 vs iLD2 versus nLD2 Figure S3a:** Induced LD2 has superior growth in *ex vivo* CSF compared with its naïve self, with higher fungal burdens after 48 hours incubation (median 5.9 ×10^4^ CFU/ml, IQR 3.6×10^4^ – 7.6×10^4^ vs 3.5 ×10^4^ CFU/ml, IQR 2.3 – 5.7 ×10^4^, P= 0.02 **b. Figure S3b: Fungal burden recovered from Galleria Figure S3b:** The 48 hour fungal burden in *Galleria* hemolymph infected with induced isolate is significantly higher than that of its naïve self (median fungal burden 9.6 ×10^6^ CFU/g body weight (inter-quartile range 6.9 ×10^6^ - 1.4×10^7^) versus 5.3 ×10^6^ CFU/g body weight (IQR 4.0 ×10^6^ - 6.7 ×10^6^, P=0.025),

**Supplementary Figure S4:**
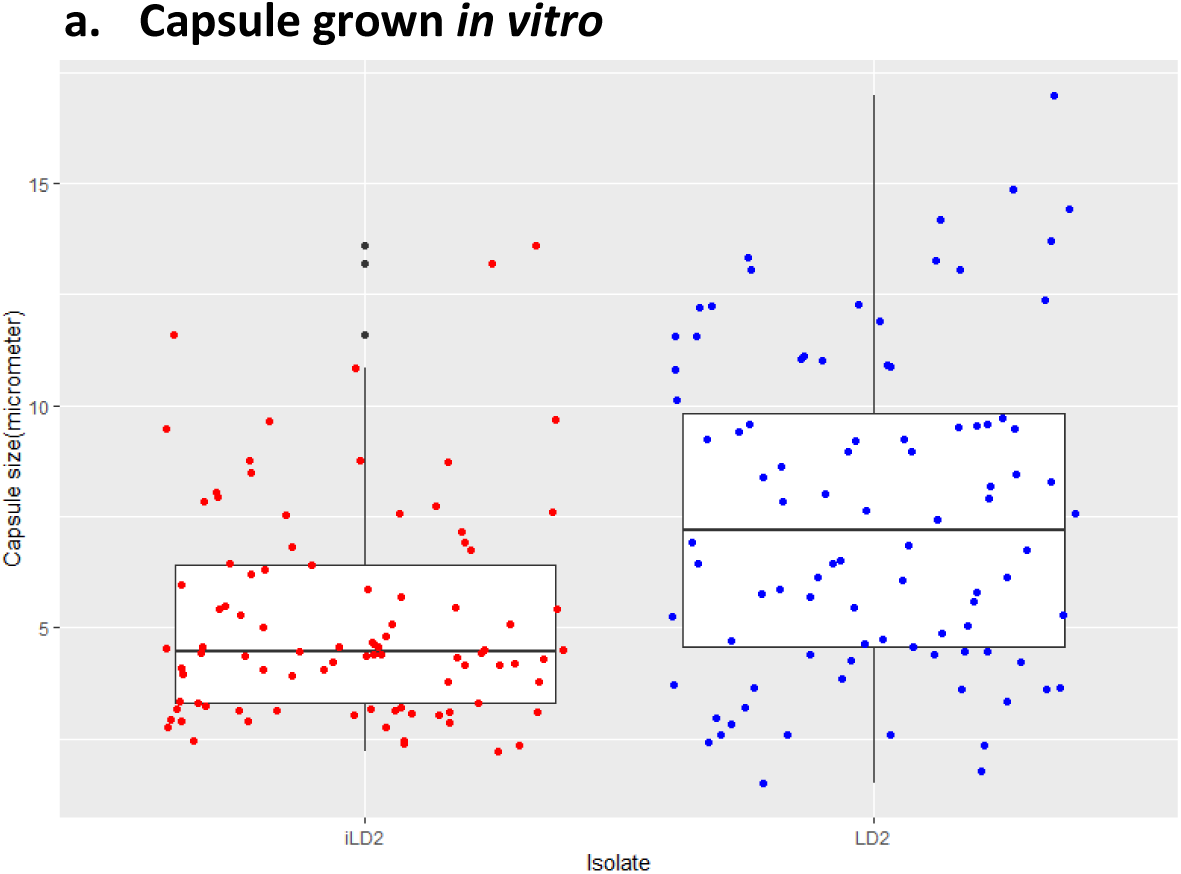
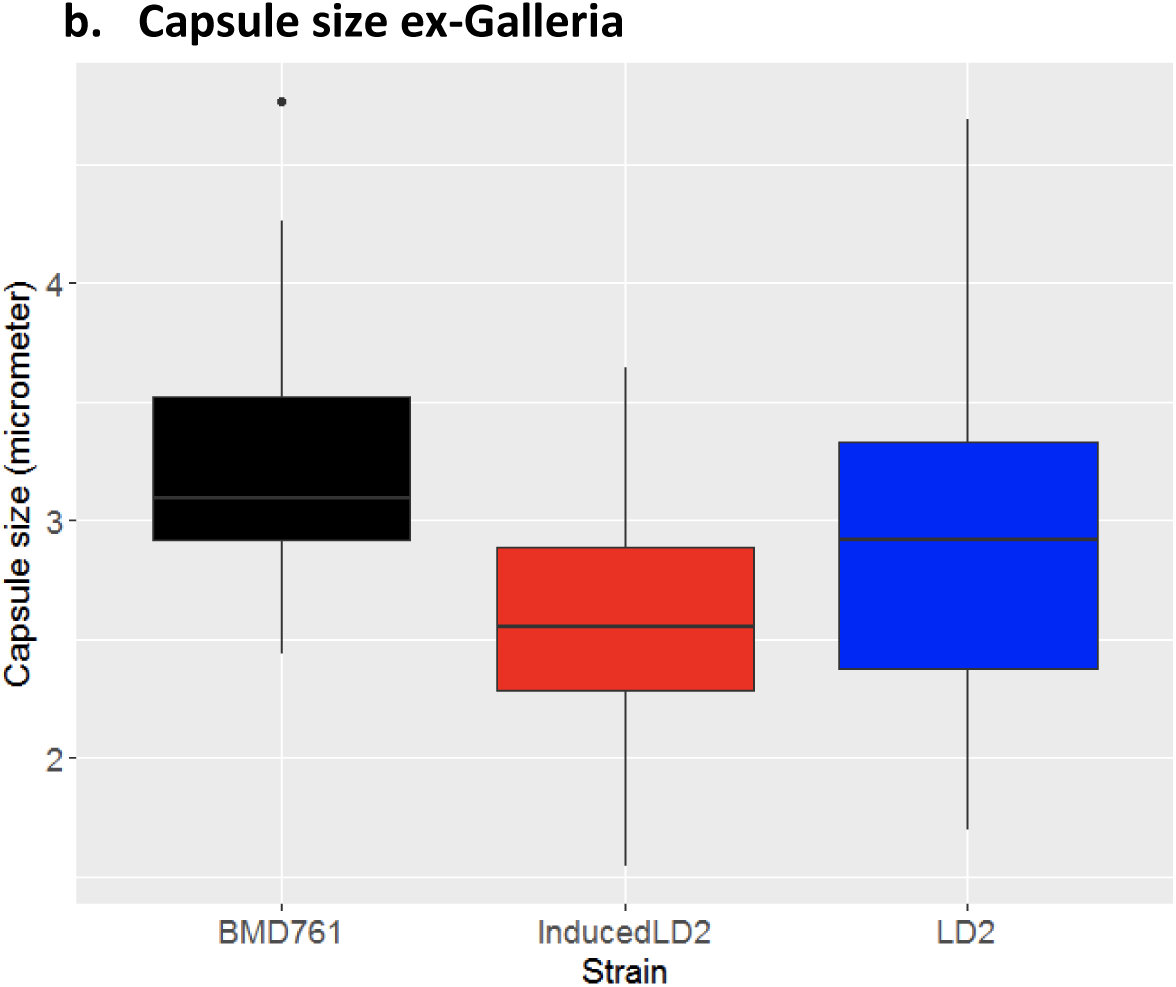
Capsule size of induced LD2 compared with naïve LD2. **a. Capsule grown *in vitro*** Induced LD2 isolates expressed significantly thinner capsules compared with the naïve self both when grown *in vitro* (median capsule width 4.5 µm, IQR 3.3-6.4 versus 7.2 µm, IQR 4.5 – 9.8, P<0.001 **b. Capsule size ex-Galleria** Capsular size comparison of *C. neoformans* (clinical BMD761, environmental LD2 and Induced LD2) recovered from larval hemolymph 48 hours post-inoculation. Adjusted P value = 0.03 obtained from Kruskal-Wallis test with Benjamini-Hochberg method (Induced LD2 vs LD2)

**Supplementary Figure S5:**
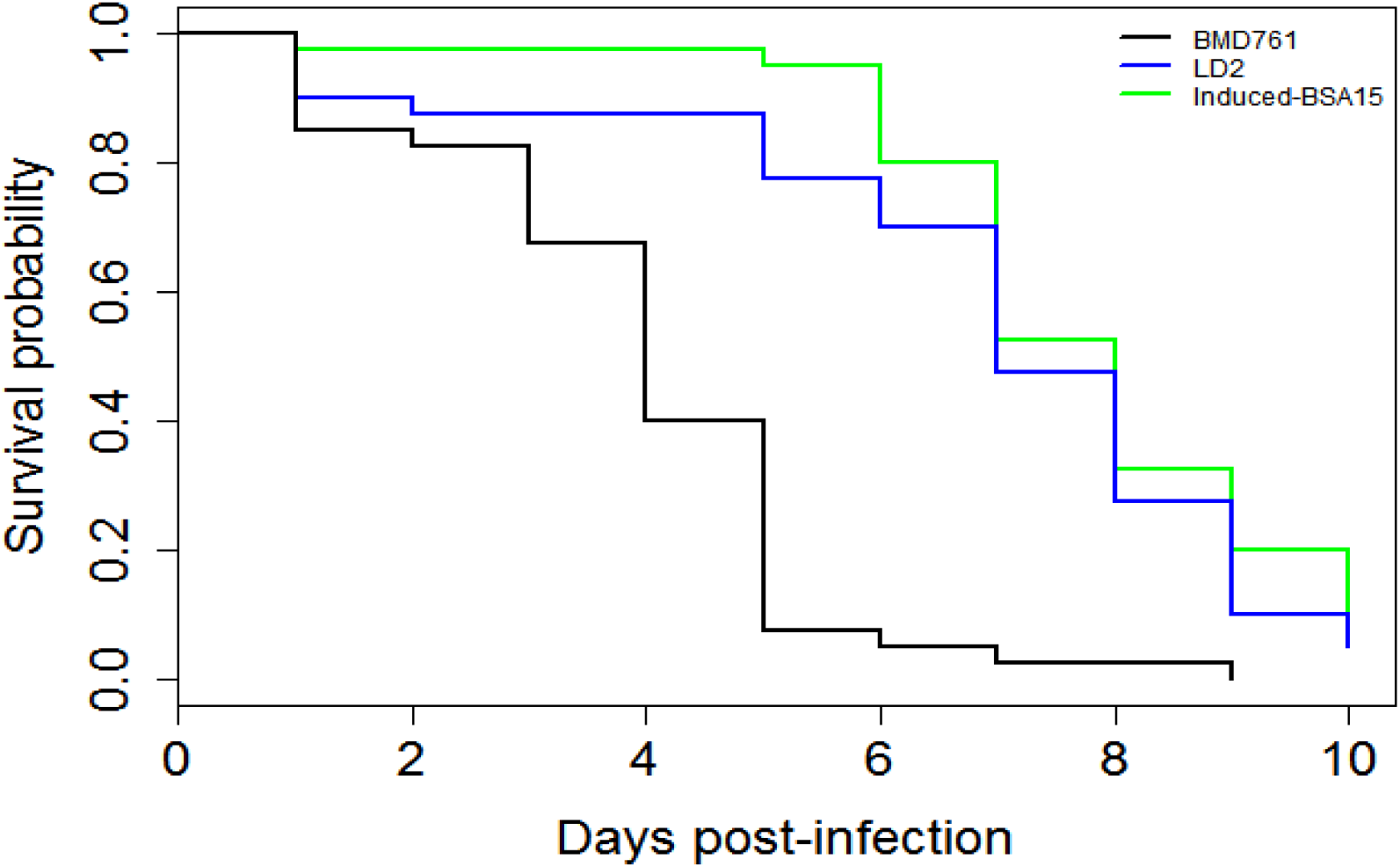
effect of bovine serum albumin on induction of environmental isolate LD2 by hypervirulent isolate BMD761. *Galleria* infection experiment following induction of naïve environmental isolate LD2 in sterile culture filtrate derived from hypervirulent VNIa-5 isolate BMD761, but following addition of bovine serum albumin. The resulting isolate is termed Induced-BSA15. Survival curves are for the infection of Galleria with the naïve environmental isolate LD2 (blue), Induced-BSA15, and BMD761. No induction of virulence in seen when BSA is added to sCF (Hazard ratio for induced-BSA15 versus LD2: HR 0.8; 95CI 0.5, 1.3, P=0.32

**Supplementary Figure S6:**
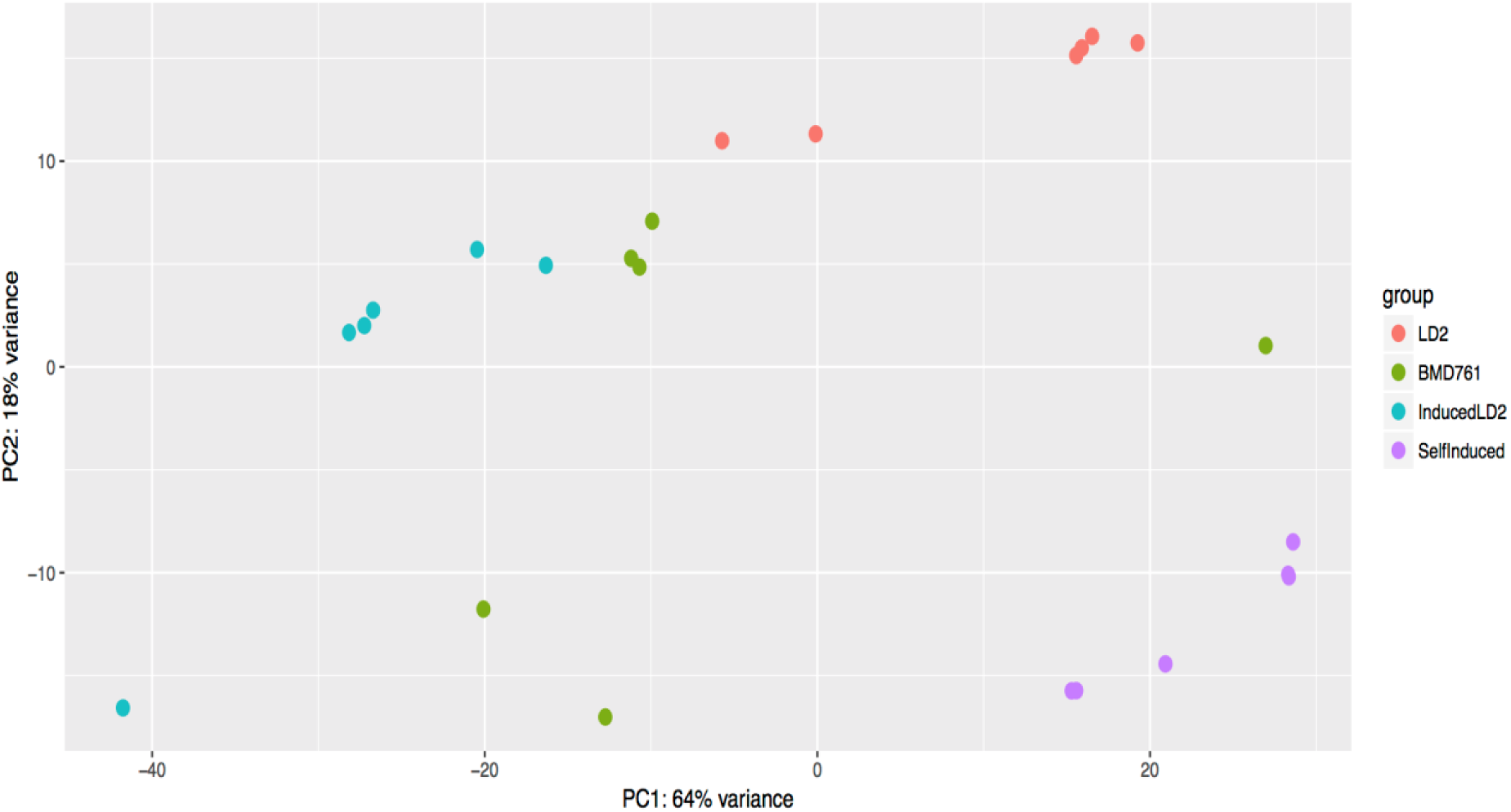
Principle components analysis of 6 biological replicates of four conditions: naïve environmental LD2, BMD761 from an HIV uninfected patient, induced LD2 cultured in the presence of supernatant from LD2, and LD2 ‘self-induced’ i.e. grown in the presence of supernatant from a different culture of LD2.

**Supplementary Figure S7:**
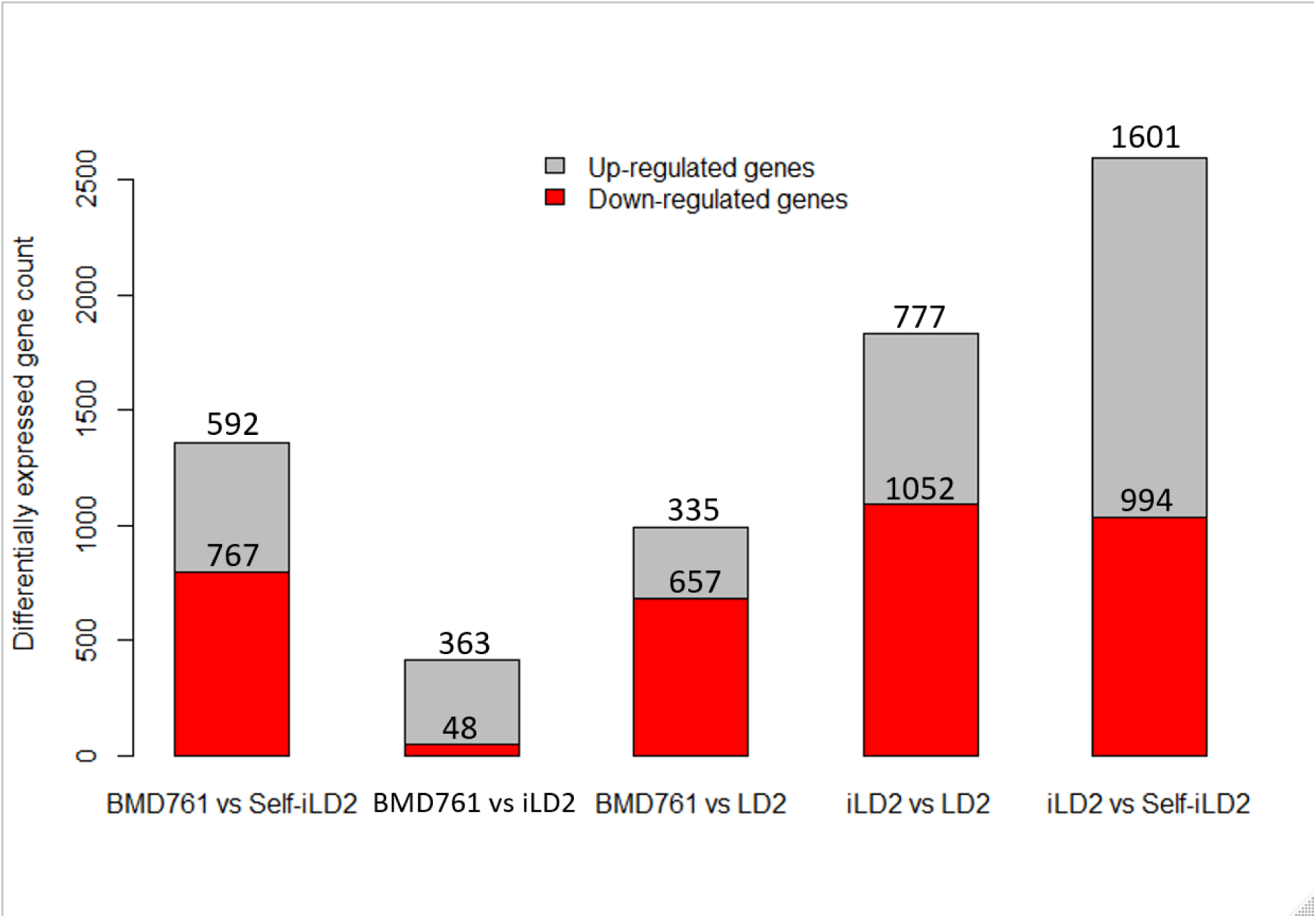
The number of differentially expressed genes between different samples (outliers included). Genes were counted as differentially expressed if they had a Benjamani-Hochberg adjusted P-value of <0.05, and log2 fold change ≥1 and ≤-1.

**Table S1:**
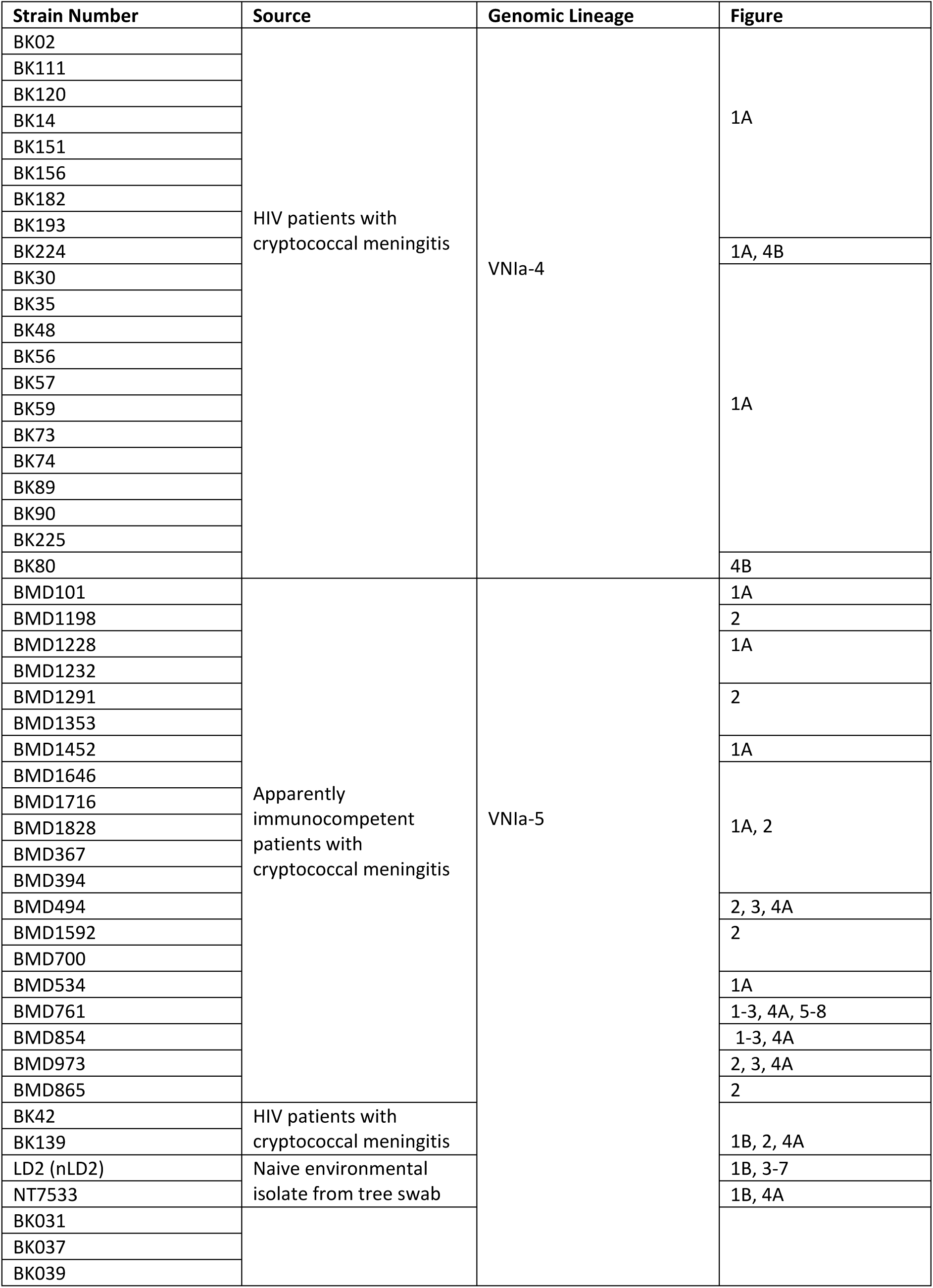
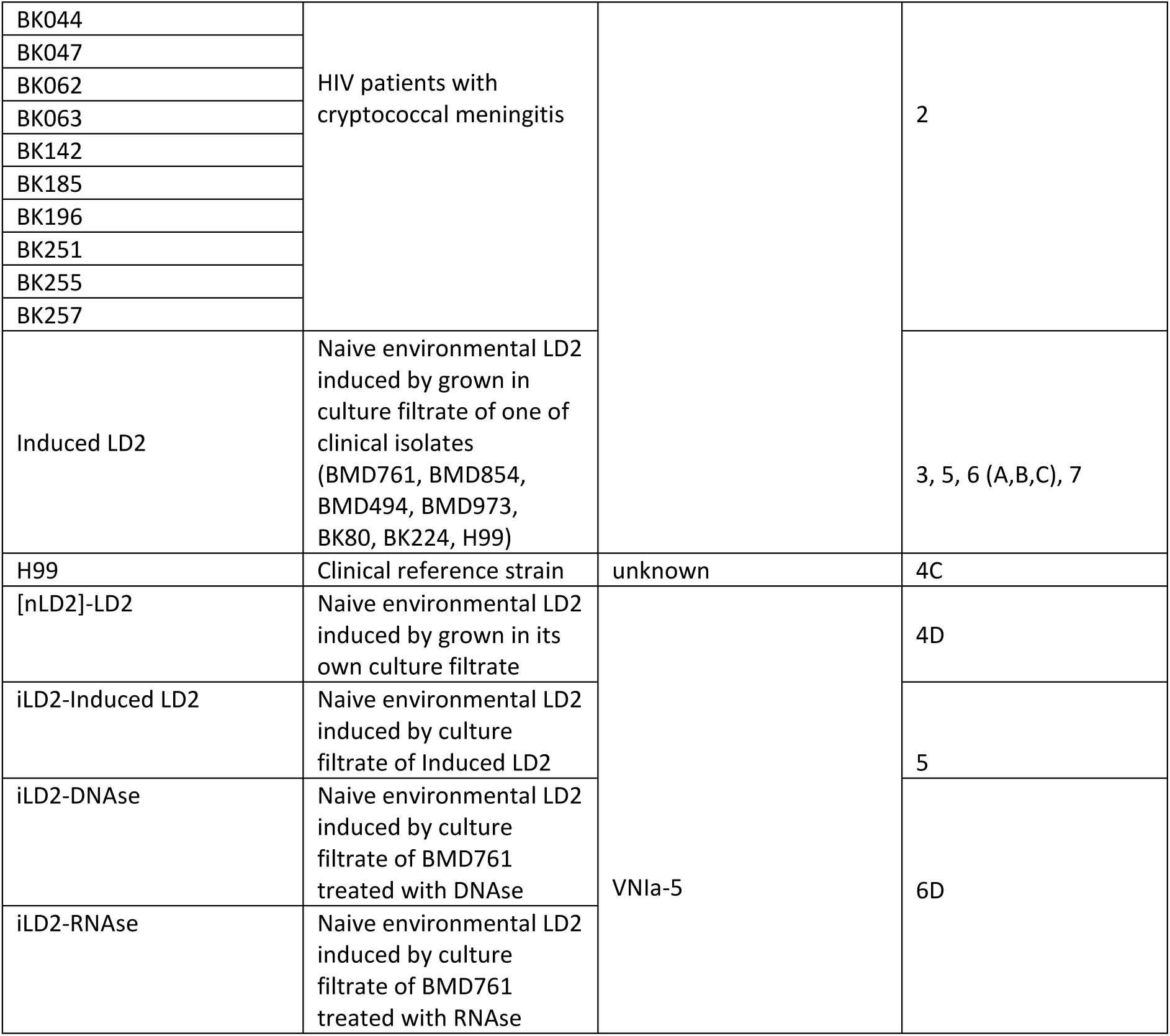
Details of strains used.

**Supplementary Table S2.**
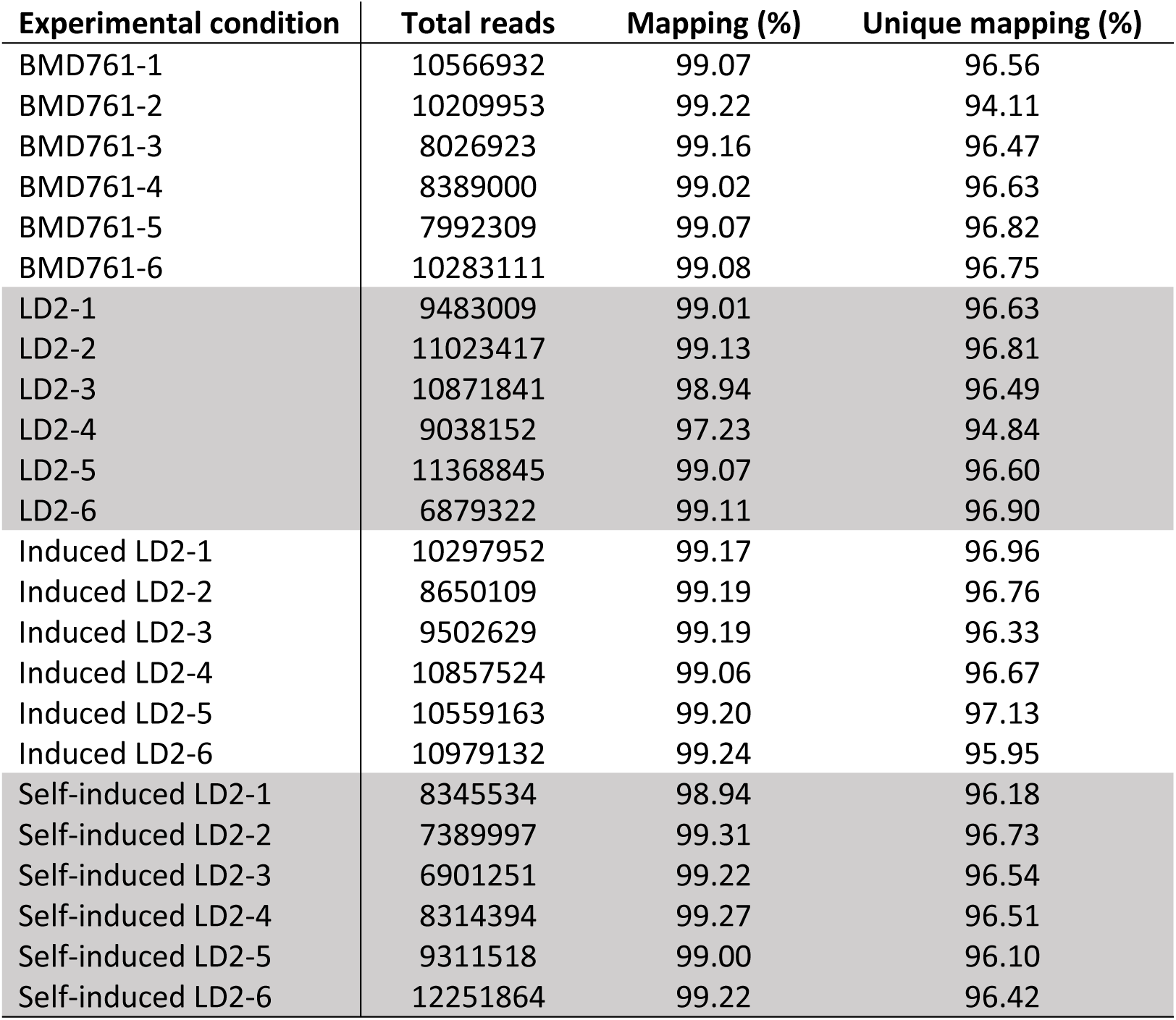
Mapping statistics for *C. neoformans* var. *grubii* RNA-seq data. Each isolate was performed in six biological replicates. Total reads: number of short read generated; Mapping: percentage of reads mapped to *C. neoformans* var. *grubii* H99 reference genome. Unique mapping: percentage of reads uniquely mapped to genomic features in the reference genome.

| Experimental condition | Total reads | Mapping (%) | Unique mapping (%) |
| --- | --- | --- | --- |
| BMD761-1 | 10566932 | 99.07 | 96.56 |
| BMD761-2 | 10209953 | 99.22 | 94.11 |
| BMD761-3 | 8026923 | 99.16 | 96.47 |
| BMD761-4 | 8389000 | 99.02 | 96.63 |
| BMD761-5 | 7992309 | 99.07 | 96.82 |
| BMD761-6 | 10283111 | 99.08 | 96.75 |
| LD2-1 | 9483009 | 99.01 | 96.63 |
| LD2-2 | 11023417 | 99.13 | 96.81 |
| LD2-3 | 10871841 | 98.94 | 96.49 |
| LD2-4 | 9038152 | 97.23 | 94.84 |
| LD2-5 | 11368845 | 99.07 | 96.60 |
| LD2-6 | 6879322 | 99.11 | 96.90 |
| Induced LD2-1 | 10297952 | 99.17 | 96.96 |
| Induced LD2-2 | 8650109 | 99.19 | 96.76 |
| Induced LD2-3 | 9502629 | 99.19 | 96.33 |
| Induced LD2-4 | 10857524 | 99.06 | 96.67 |
| Induced LD2-5 | 10559163 | 99.20 | 97.13 |
| Induced LD2-6 | 10979132 | 99.24 | 95.95 |
| Self-induced LD2-1 | 8345534 | 98.94 | 96.18 |
| Self-induced LD2-2 | 7389997 | 99.31 | 96.73 |
| Self-induced LD2-3 | 6901251 | 99.22 | 96.54 |
| Self-induced LD2-4 | 8314394 | 99.27 | 96.51 |
| Self-induced LD2-5 | 9311518 | 99.00 | 96.10 |
| Self-induced LD2-6 | 12251864 | 99.22 | 96.42 |

**Table S3:** Regulation of virulence-associated genes in BMD761 and Induced LD2 strains relative to naive LD2. Null means there is no function described or no ortholog for the gene in question.

| GeneID | Product Description | Name | log2FoldChange |
| --- | --- | --- | --- |
| CNAG_05472 | Meiotic recombination protein | SPO11 | -2.1 |
| CNAG_06574 | Secreted antiphagocytic protein | APP1 | -2.1 |
| CNAG_05015 | Catalase 4 | CAT4 | -1.7 |
| CNAG_03012 | Quorum sensing protein 1 | QSP1 | -1.3 |
| CNAG_04217 | Phosphoenolpyruvate carboxykinase (ATP) | PCK1 | -1.3 |
| CNAG_05181 | White-collar transcription factor; blue light photoresponsive gene | BWC1 | -1.0 |
| CNAG_06500 | Arginine metabolism transcriptional control protein | ARG1 | 1.0 |
| CNAG_02885 | Capsule-associated protein | CAP64 | 1.1 |
| CNAG_02729 | Hypothetical protein | FRR1 | 1.2 |
| CNAG_03263 | Translation elongation factor Tu | TUF1 | 1.2 |
| CNAG_02531 | Mitogen activated protein kinase | CPK2 | 1.2 |
| CNAG_05998 | Rho-GTPase | RAC2 | 1.4 |
| CNAG_01172 | Parallel beta-helix repeat protein | PBX1 | 1.5 |
| CNAG_05274 | STE/STE20/YSK protein kinase | null | 1.6 |
| CNAG_00418 | S-adenosylmethionine synthase | ILV2 | 1.6 |
| CNAG_02036 | Putative sugar transporter | CAS4 | 1.7 |
| CNAG_01890 | Methionine synthase | MET6 | 1.7 |
| CNAG_05968 | Rho-like GTPase | CDC420 | 1.7 |
| CNAG_06231 | Large subunit ribosomal protein L13 | null | 2.1 |
| CNAG_00779 | Large subunit ribosomal protein L27e | RPL27 | 2.4 |
| CNAG_06487 | Putative chitin synthase | CHS6 | 4.6 |
| CNAG_01907 | PLK/PLK1 protein kinase | null | 5.3 |
| CNAG_05565 | Hypothetical protein | null | 5.7 |

## References

1. Rajasingham R, Smith RM, Park BJ, Jarvis JN, Govender NP, Chiller TM, Denning DW, Loyse A, Boulware DR. Global burden of disease of HIV-associated cryptococcal meningitis: an updated analysis. Lancet Infect Dis. 2017;17(8):873–881.

2. Ashton PM, Thanh LT, Trieu PH, Van Anh D, Trinh NM, Beardsley J, Kibengo F, Chierakul W, Dance DAB, Rattanavong S, Davong V, Hung LQ, Chau NVV, Tung NLN, Chan AK, Thwaites GE, Lalloo DG, Anscombe C, Nhat LTH, Perfect J, Dougan G, Baker S, Harris S, Day JN. Three phylogenetic groups have driven the recent population expansion of Cryptococcus neoformans. Nat Commun. 2019;10(1):2035.

3. Day JN, Qihui S, Thanh LT, Trieu PH, Van AD, Thu NH, Chau TTH, Lan NPH, Chau NVV, Ashton PM, Thwaites GE, Boni MF, Wolbers M, Nagarajan N, Tan PBO, Baker S. Comparative genomics of Cryptococcus neoformans var. grubii associated with meningitis in HIV infected and uninfected patients in Vietnam. PLoS Negl Trop Dis. 2017;11(6):e0005628.

4. Thanh LT, Phan TH, Rattanavong S, Nguyen TM, Duong AV, Dacon C, Hoang TN, Nguyen LPH, Tran CTH, Davong V, Nguyen CVV, Thwaites GE, Boni MF, Dance D, Ashton PM, Day JN. Multilocus sequence typing of Cryptococcus neoformans var. grubii from Laos in a regional and global context. Med Mycol. 2018.

5. Chen J, Varma A, Diaz MR, Litvintseva AP, Wollenberg KK, Kwon-Chung KJ. Cryptococcus neoformans strains and infection in apparently immunocompetent patients, China. Emerg Infect Dis. 2008;14(5):755–762.

6. Chen YC, Chang SC, Shih CC, Hung CC, Luhbd KT, Pan YS, Hsieh WC. Clinical features and in vitro susceptibilities of two varieties of Cryptococcus neoformans in Taiwan. Diagn Microbiol Infect Dis. 2000;36(3):175–183.

7. Choi YH, Ngamskulrungroj P, Varma A, Sionov E, Hwang SM, Carriconde F, Meyer W, Litvintseva AP, Lee WG, Shin JH, Kim EC, Lee KW, Choi TY, Lee YS, Kwon-Chung KJ. Prevalence of the VNIc genotype of Cryptococcus neoformans in non-HIV-associated cryptococcosis in the Republic of Korea. FEMS Yeast Res. 2010;10(6):769–778.

8. Feng X, Yao Z, Ren D, Liao W, Wu J. Genotype and mating type analysis of Cryptococcus neoformans and Cryptococcus gattii isolates from China that mainly originated from non-HIV-infected patients. FEMS Yeast Res. 2008.

9. Chau TT, Mai NH, Phu NH, Nghia HD, Chuong LV, Sinh DX, Duong VA, Diep PT, Campbell JI, Baker S, Hien TT, Lalloo DG, Farrar JJ, Day JN. A prospective descriptive study of cryptococcal meningitis in HIV uninfected patients in Vietnam - high prevalence of Cryptococcus neoformans var grubii in the absence of underlying disease. BMC Infect Dis. 2010;10:199.

10. Casadevall A, Perfect JR. Cryptococcus neoformans. 1 ed. Washington: American Society for Microbiology Press; 1998.

11. Bartlett KH, Kidd SE, Kronstad JW. The Emergence of Cryptococcus gattii in British Columbia and the Pacific Northwest. Curr Infect Dis Rep. 2008;10(1):58–65.

12. Day JN, Hoang TN, Duong AV, Hong CT, Diep PT, Campbell JI, Sieu TP, Hien TT, Bui T, Boni MF, Lalloo DG, Carter D, Baker S, Farrar JJ. Most Cases of Cryptococcal Meningitis in HIV-Uninfected Patients in Vietnam Are Due to a Distinct Amplified Fragment Length Polymorphism-Defined Cluster of Cryptococcus neoformans var. grubii VN1. J Clin Microbiol. 2011;49(2):658–664.

13. Bielska E, Sisquella MA, Aldeieg M, Birch C, O’Donoghue EJ, May RC. Pathogen-derived extracellular vesicles mediate virulence in the fatal human pathogen Cryptococcus gattii. Nat Commun. 2018;9(1):1556.

14. Rodrigues ML, Nakayasu ES, Oliveira DL, Nimrichter L, Nosanchuk JD, Almeida IC, Casadevall A. Extracellular vesicles produced by Cryptococcus neoformans contain protein components associated with virulence. Eukaryot Cell. 2008;7(1):58–67.

15. Rodrigues ML, Nimrichter L, Oliveira DL, Frases S, Miranda K, Zaragoza O, Alvarez M, Nakouzi A, Feldmesser M, Casadevall A. Vesicular polysaccharide export in Cryptococcus neoformans is a eukaryotic solution to the problem of fungal trans-cell wall transport. Eukaryot Cell. 2007;6(1):48–59.

16. Wolf JM, Rivera J, Casadevall A. Serum albumin disrupts Cryptococcus neoformans and Bacillus anthracis extracellular vesicles. Cell Microbiol. 2012;14(5):762–773.

17. Sayers S, Li L, Ong E, Deng S, Fu G, Lin Y, Yang B, Zhang S, Fa Z, Zhao B, Xiang Z, Li Y, Zhao XM, Olszewski MA, Chen L, He Y. Victors: a web-based knowledge base of virulence factors in human and animal pathogens. Nucleic Acids Res. 2019;47(D1):D693–D700.

18. Homer CM, Summers DK, Goranov AI, Clarke SC, Wiesner DL, Diedrich JK, Moresco JJ, Toffaletti D, Upadhya R, Caradonna I, Petnic S, Pessino V, Cuomo CA, Lodge JK, Perfect J, Yates JR, 3rd, Nielsen K, Craik CS, Madhani HD. Intracellular Action of a Secreted Peptide Required for Fungal Virulence. Cell Host Microbe. 2016;19(6):849–864.

19. Edskes HK, Khamar HJ, Winchester CL, Greenler AJ, Zhou A, McGlinchey RP, Gorkovskiy A, Wickner RB. Sporadic distribution of prion-forming ability of Sup35p from yeasts and fungi. Genetics. 2014;198(2):605–616.

20. Jurka J, Kapitonov VV, Pavlicek A, Klonowski P, Kohany O, Walichiewicz J. Repbase Update, a database of eukaryotic repetitive elements. Cytogenet Genome Res. 2005;110(1-4):462–467.

21. Albuquerque PC, Nakayasu ES, Rodrigues ML, Frases S, Casadevall A, Zancope-Oliveira RM, Almeida IC, Nosanchuk JD. Vesicular transport in Histoplasma capsulatum: an effective mechanism for trans-cell wall transfer of proteins and lipids in ascomycetes. Cell Microbiol. 2008;10(8):1695–1710.

22. Rodrigues ML, Nimrichter L, Oliveira DL, Nosanchuk JD, Casadevall A. Vesicular Trans-Cell Wall Transport in Fungi: A Mechanism for the Delivery of Virulence-Associated Macromolecules? Lipid Insights. 2008;2:27–40.

23. Rodrigues ML, Casadevall A. A two-way road: novel roles for fungal extracellular vesicles. Mol Microbiol. 2018;110(1):11–15.

24. Ofir-Birin Y, Regev-Rudzki N. Extracellular vesicles in parasite survival. Science. 2019;363(6429):817–818.

25. Konoshenko MY, Lekchnov EA, Vlassov AV, Laktionov PP. Isolation of Extracellular Vesicles: General Methodologies and Latest Trends. Biomed Res Int. 2018;2018:8545347.

26. Casadevall A. Amoeba provide insight into the origin of virulence in pathogenic fungi. Adv Exp Med Biol. 2012;710:1–10.

27. Casadevall A, Fu MS, Guimaraes AJ, Albuquerque P. The ‘Amoeboid Predator-Fungal Animal Virulence’ Hypothesis. J Fungi (Basel). 2019;5(1).

28. Coelho C, Bocca AL, Casadevall A. The tools for virulence of Cryptococcus neoformans. Adv Appl Microbiol. 2014;87:1–41.

29. Derengowski Lda S, Paes HC, Albuquerque P, Tavares AH, Fernandes L, Silva-Pereira I, Casadevall A. The transcriptional response of Cryptococcus neoformans to ingestion by Acanthamoeba castellanii and macrophages provides insights into the evolutionary adaptation to the mammalian host. Eukaryot Cell. 2013;12(5):761–774.

30. Perfect JR, Dismukes WE, Dromer F, Goldman DL, Graybill JR, Hamill RJ, Harrison TS, Larsen RA, Lortholary O, Nguyen MH, Pappas PG, Powderly WG, Singh N, Sobel JD, Sorrell TC. Clinical Practice Guidelines for the Management of Cryptococcal Disease: 2010 Update by the Infectious Diseases Society of America. Clin Infect Dis. 2010;50(3):291–322.

31. Chen Y, Toffaletti DL, Tenor JL, Litvintseva AP, Fang C, Mitchell TG, McDonald TR, Nielsen K, Boulware DR, Bicanic T, Perfect JR. The Cryptococcus neoformans transcriptome at the site of human meningitis. mBio. 2014;5(1):e01087–01013.

32. Blackburn SL, Grande AW, Swisher CB, Hauck EF, Jagadeesan B, Provencio JJ. Prospective Trial of Cerebrospinal Fluid Filtration After Aneurysmal Subarachnoid Hemorrhage via Lumbar Catheter (PILLAR). Stroke. 2019;50(9):2558–2561.

33. Blackburn SL, Swisher CB, Grande AW, Rubi A, Verbick LZ, McCabe A, Lad SP. Novel Dual Lumen Catheter and Filtration Device for Removal of Subarachnoid hemorrhage: First Case Report. Oper Neurosurg (Hagerstown). 2019;16(5):E148–E153.

34. Smilnak GJ, Charalambous LT, Cutshaw D, Premji AM, Giamberardino CD, Ballard CG, Bartuska AP, Ejikeme TU, Sheng H, Verbick LZ, Hedstrom BA, Pagadala PC, McCabe AR, Perfect JR, Lad SP. Novel Treatment of Cryptococcal Meningitis via Neurapheresis Therapy. J Infect Dis. 2018;218(7):1147–1154.

35. Beardsley J, Wolbers M, Kibengo FM, Ggayi AB, Kamali A, Cuc NT, Binh TQ, Chau NV, Farrar J, Merson L, Phuong L, Thwaites G, Van Kinh N, Thuy PT, Chierakul W, Siriboon S, Thiansukhon E, Onsanit S, Supphamongkholchaikul W, Chan AK, Heyderman R, Mwinjiwa E, van Oosterhout JJ, Imran D, Basri H, Mayxay M, Dance D, Phimmasone P, Rattanavong S, Lalloo DG, Day JN, CryptoDex I. Adjunctive Dexamethasone in HIV-Associated Cryptococcal Meningitis. N Engl J Med. 2016;374(6):542–554.

36. Day JN, Chau TT, Wolbers M, Mai PP, Dung NT, Mai NH, Phu NH, Nghia HD, Phong ND, Thai CQ, Thai le H, Chuong LV, Sinh DX, Duong VA, Hoang TN, Diep PT, Campbell JI, Sieu TP, Baker SG, Chau NV, Hien TT, Lalloo DG, Farrar JJ. Combination antifungal therapy for cryptococcal meningitis. N Engl J Med. 2013;368(14):1291–1302.

37. Lee A, Toffaletti DL, Tenor J, Soderblom EJ, Thompson JW, Moseley MA, Price M, Perfect JR. Survival defects of Cryptococcus neoformans mutants exposed to human cerebrospinal fluid result in attenuated virulence in an experimental model of meningitis. Infect Immun. 2010;78(10):4213–4225.

38. Zaragoza O, Casadevall A. Experimental modulation of capsule size in Cryptococcus neoformans. Biol Proced Online. 2004;6:10–15.

39. Sabiiti W, Robertson E, Beale MA, Johnston SA, Brouwer AE, Loyse A, Jarvis JN, Gilbert AS, Fisher MC, Harrison TS, May RC, Bicanic T. Efficient phagocytosis and laccase activity affect the outcome of HIV-associated cryptococcosis. J Clin Invest. 2014;124(5):2000–2008.

40. Chen SC, Muller M, Zhou JZ, Wright LC, Sorrell TC. Phospholipase activity in Cryptococcus neoformans: a new virulence factor? J Infect Dis. 1997;175(2):414–420.

41. Liao Y, Smyth GK, Shi W. featureCounts: an efficient general purpose program for assigning sequence reads to genomic features. Bioinformatics. 2014;30(7):923–930.

42. Love MI, Huber W, Anders S. Moderated estimation of fold change and dispersion for RNA-seq data with DESeq2. Genome Biol. 2014;15(12):550.

43. Love MI, Anders S, Kim V, Huber W. RNA-Seq workflow: gene-level exploratory analysis and differential expression. F1000Res. 2015;4:1070.

44. R Core Team. R: A language and environment for statistical computing. 2018; http://www.R-project.org. Accessed 01 January 2020.

45. Kassambara A, Kosinskis M, Przemyslaw B, Scheipl F. survminer: Survival Analysis and Visualization. 2016. https://rpkgs.datanovia.com/survminer/index.html

46. Wickham H. ggplot2: Elegant Graphics for Data Analysis.: Springer-Verlag New York.; 2009.

